# SP2 impedes the pluripotent-to-totipotent state transition in an RA-dependent manner

**DOI:** 10.64898/2026.07.23.740258

**Authors:** Siling Hu, Pinghui Zhu, He Liu, Huan Chen, Shunlan Xu, Enjing Guo, Baomei Cai, Xiongzhi Quan, Duanqing Pei, Jiekai Chen, Haoyu Wu, Shangtao Cao

## Abstract

The pluripotency-to-totipotency transition represents a transient developmental reversal that remains not fully understood. Although chemical approaches can induce a 2-cell (2C)-like state in mouse embryonic stem cells (mESCs), how cells choose between totipotency acquisition and alternative developmental fates remains unclear. Here, we established a chemical induction system that efficiently drives mESCs toward the 2C-like state. We found that mESCs under induction treatment followed distinct trajectories, with a subset activating developmental programs rather than acquiring 2C-like features. SP2 was preferentially expressed along this developmental trajectory and progressively reduced during successful 2C-like induction. Loss of SP2 enhanced 2C-like cell formation, whereas SP2 overexpression impaired the transition. Transcriptomic analysis of SP2-deficient cells showed reduced expression of developmental genes, including *Hox* family genes, accompanied by increased expression of 2C-associated genes. Consistent with this finding, SP2 occupied the promoters of *Hox* family genes in the reprogrammed cells. Notably, the effect of SP2 loss was particularly evident upon activation of retinoic acid (RA) signaling, linking SP2-dependent developmental gene activation to the RA-responsive branch of the transition. Altogether, our findings identify SP2 function as a barrier that limits acquisition of the 2C-like state by promoting a competing developmental program, highlighting the importance of coordinated extracellular signaling and transcriptional regulation in directing cell fate transitions.

## Introduction

In mice, the zygote and early cleavage-stage blastomeres possess totipotency, the capacity to generate an entire organism, including all cell types from both embryonic and extraembryonic lineages^1, 2^. At the transition from morula to blastocyst, cells undergo the first fate decision, differentiating into two distinct lineages: the trophectoderm and the inner cell mass (ICM)^3^. Mouse embryonic stem cells (mESCs) derived from ICM are pluripotent with a restricted differentiation potential that enables the generation of all somatic cell types but not extraembryonic tissues^4^. Although transient *in vivo*, totipotent cells provide an important window to understand the earliest cell fate decisions during mammalian development. Previous studies have identified a rare subpopulation within mESC cultures, referred to as two-cell-like cells (2CLCs), that displays transcriptional and epigenetic features reminiscent of two-cell (2C) stage embryos^5, 6^, providing a perfect model for investigating the transition from pluripotency to totipotency.

Previous studies have identified multiple regulators involved in this process, including transcription factors (TFs)^7, 8^, epigenetic modifiers^9–12^, chromatin regulators^13, 14^, and extracellular signaling pathways that regulate the acquisition of 2C-like features. Retinoic acid (RA) signaling, which plays crucial roles during embryonic patterning and lineage specification, has recently been implicated in promoting pluripotency-to-totipotency transition. Several RA agonists, including ATRA, TTNPB, CD1530, and AM580, have been utilized in 2CLC induction systems with expanded developmental potential^15–18^. However, RA signaling is also a major trigger of developmental differentiation, largely through activation of downstream developmental genes. How these distinct cell fates, namely reverting toward a totipotent-like state or entering differentiation programs, are regulated upon the same developmental signal remains largely unexplored.

In this study, we established a highly efficient chemical approach for inducing 2C-like cells from mESCs and used this system to investigate the mechanisms underlying RA-mediated cell fate transition. We found that AM580, an RA receptor agonist, strongly promotes the acquisition of totipotent-like features while simultaneously inducing a subset of mESCs toward differentiation. Single-cell transcriptomic analysis revealed distinct cell populations following AM580 treatment, corresponding to successful reprogramming and differentiation-associated states. Motif enrichment analysis revealed significant enrichment of SP2 motifs in the promoters of marker genes specific to the differentiation-associated cell population.

Mechanistically, SP2 directly binds and promotes the expression of developmental genes, thereby favoring differentiation and limiting the conversion toward a totipotent-like state. Removal of SP2 significantly increases the proportion of cells acquiring 2C-like features under RA activation. Together, our findings reveal a previously unrecognized role of SP2 in controlling the balance between totipotency acquisition and developmental differentiation in an RA-dependent manner, providing insights into how signaling pathways regulate early cell fate transitions.

## Results

### A robust and fast 2C-like reprogramming approach with 7 chemical molecules

Previous studies have shown that treatment with small molecules and signal agonists can increase the frequency of 2C-like cells in mESC culture. Based on these findings, we reasoned that pharmacological modulation of pathways associated with pluripotency exit, chromatin regulation, and early embryonic transcriptional programs could facilitate the conversion of mESCs toward a 2C-like state. We therefore established a defined chemical induction system incorporating a DOT1L inhibitor (EPZ5676), an HDAC inhibitor (VPA), and the RA receptor agonist (AM580), together with additional compounds reported to promote cell fate reprogramming^19–22^. We refer to this defined culture condition as “Chemical induction medium (CIM)” (**Fig. 1a**). To monitor the acquisition of 2C-like features, we utilized an MERVL-tdTomato reporter mESC line and quantified the percentage of MERVL-tdTomato^+^ cells with Fluorescence-activated cell sorting (FACS). Treatment with the CIM induced a rapid increase in MERVL-tdTomato^+^ cells, reaching a maximum of more than 70% within 36 h (**Fig. 1a, b**). Thus, this system enabled efficient conversion of mESCs into an MERVL-positive population within a relatively short period. Core pluripotency genes such as *Oct4*, *Nanog*, and *Sox2*, gradually decreased during the induction, whereas the totipotency genes including *Zscan4*, *Tcstv1*, and *Tcstv3*, increased progressively (**Supplementary Fig. S1a**). These changes were consistent with a transition from the conventional pluripotent state toward a 2C-like state.

**Fig. 1.**
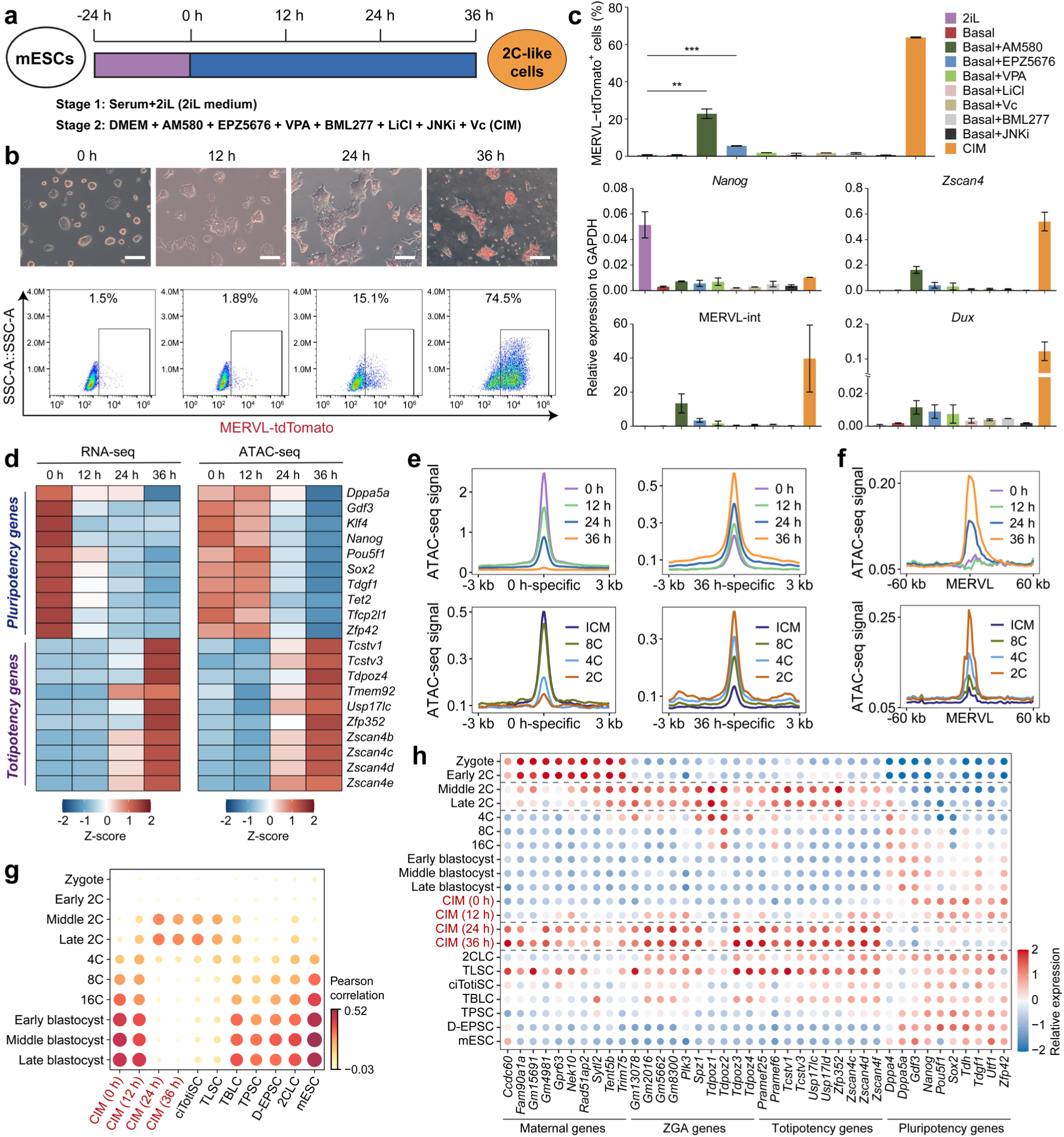
Induction of mESCs into 2C-like cells using a chemical cocktail. **a,** Schematic diagram for the induction of 2C-like cells from mESCs. Cells were first treated in 2i/LIF medium for at least 24h (stage 1), and then switch to chemical induction medium (CIM) containing chemical molecules listed below (stage2). **b,** Merged cell images showing the morphological changes of mESCs during CIM treatment at 0 h, 12 h, 24 h and 36 h with the overlapping of tdTomato signal on the top panel, and the FACS analysis showing the percentage of MERVL-tdTomato^+^ cells on the bottom panel. Scale bars = 250 μm. **c,** Upper panel: Bar plot showing the percentage of MERVL-tdTomato^+^ cells under each culture condition. Lower panel: Bar plots showing the expression of *Nanog*, *Zscan4*, MERVL-int, and *Dux* under each culture condition. Data are mean ± s.d., *n* = 2 independent experiments. \*\**P*<0.01, \*\*\**P*<0.001; two-tailed unpaired Student’s *t*-test. 2iL, 2iL medium; Basal: basal medium. **d**, Heatmap showing the relative expression and chromatin accessibility around the transcription start site (TSS) of representative totipotency and pluripotency genes in chemical reprogramming cells at 0 h, 12 h, 24 h and 36 h. **e**, Average profiles showing normalized ATAC-seq signals of CIM-treated cells (0 h, 12 h, 24 h and 36 h), early mouse embryos (from 2C to ICM) centered around peaks gained or lost at 36 h relative to 0 h. **f**, Average profiles showing normalized ATAC-seq signals of CIM-treated cells (0 h, 12 h, 24 h and 36 h), early mouse embryos (from 2C to ICM) centered around the whole-genome-wide MERVLs (MT2-Mm and MERVL-int). **g**, Pearson correlation analysis of bulk RNA-seq data from TLSCs, TPSCs, D-EPSCs, 2CLCs, and mESCs, together with scRNA-seq data from CIM-treated cells (0 h, 12 h, 24 h and 36 h), ciTotiSCs, TBLCs and various stages of early mouse embryos (zygote to blastocyst), based on curated totipotency and pluripotency gene sets. **h**, Relative expression of maternal, ZGA, totipotency and pluripotency genes in CIM-treated cells (0 h, 12 h, 24 h and 36 h), 2CLC, TLSCs, ciTotiSCs, TBLCs, TPSCs, D-EPSCs, mESCs detected by bulk RNA-seq, and in mouse embryos (zygote to blastocyst) detected by scRNA-seq.

To further elucidate the contribution of each small molecule in the cocktail, we performed systematic omission experiments by removing one compound at a time. The loss of any single component reduced the efficiency of mESCs conversion into 2C-like cells, with AM580, EPZ5676, and VPA exerting the most critical effects, together with attenuated induction of totipotency genes *Zscan4*, *Dux*, and MERVL-int (**Supplementary Fig. S1b, c**). Consistent with these findings, we found that single addition of each small molecule also promoted, to varying degrees, the transition of mESCs into MERVL-tdTomato^+^ cells, with AM580 exhibiting the most pronounced effect (**Fig. 1c; Supplementary Fig. S1d**). These results further demonstrate the importance of RA signaling activation during the 2C-like induction.

We next assessed the robustness of this system across different genetic backgrounds. MERVL-tdTomato reporter mESC lines derived from C57BL/6 and 129 strains were cultured in CIM. FACS analysis revealed reprogramming efficiencies of approximately 50% and 30% MERVL-tdTomato^+^ cells in the C57BL/6 and 129 backgrounds (**Supplementary Fig. S1e**), indicating that the system could induce 2C-like cells in mESCs of different genetic backgrounds. In contrast, *Dux* knockout (KO) mESCs failed to produce MERVL-tdTomato^+^ cells under the same conditions (**Supplementary Fig. S1f)**, consistent with the requirement of DUX for MERVL activation during 2C-like reprogramming.

Altogether, these results establish a defined chemical system that rapidly and efficiently induces a 2C-like state in mESCs. The contribution of individual components further indicates that RA signaling activation, DOT1L inhibition, and HDAC inhibition are important for this conversion, with AM580 showing a prominent effect on the induction of 2C-associated features.

### CIM-induced cells possess transcriptomic and epigenomic features similar to 2-cell blastomeres

We next characterized the molecular changes of these cells accompanying the transition toward the 2C-like state. Bulk RNA-seq and ATAC-seq were performed at 0, 12, 24 and 36 h after induction, together with scRNA-seq to resolve the heterogeneity of the induced population. RNA-seq analysis showed a progressive reduction in the expression of core pluripotency genes and a concomitant increase in 2C-associated genes during induction (**Fig. 1d; Supplementary Fig. S2a**). These transcriptional changes were accompanied by corresponding changes in chromatin accessibility at their regulatory regions (**Fig. 1d; Supplementary Fig. S2b**). Gene set enrichment analysis (GSEA) using established pluripotency- and totipotency-associated gene sets further supported a gradual shift from the pluripotent toward the totipotent state (**Supplementary Fig. S2c**). Changes in chromatin accessibility were also evident at loci associated with the 2C state. Compared with the starting mESCs, 2C embryo-associated loci became more accessible during induction, whereas accessibility at ICM-associated loci was reduced (**Fig. 1e, f; Supplementary Fig. S2d**). Consistent with these changes, motifs of pluripotency-associated transcription factors, including OCT4, SOX2 and KLF4, were enriched in ATAC-seq peaks specific to the 0 h state. In contrast, peaks preferentially detected at 36 h were enriched for binding motifs of 2C-associated transcription factors (**Supplementary Fig. S2d, e**). Together, these data indicate that the transition induced by the chemical cocktail involves coordinated changes in both gene expression and chromatin accessibility.

To further characterize CIM-induced cells, we performed Pearson correlation analysis to compare our cells and other 2C-like cells induced by various published strategies^16, 17, 23, 24^ with early mouse embryos^25^. We found that cells collected at 24 and 36 h displayed the highest similarity to both middle- and late-2C embryos among the populations examined, with a transcriptional relationship comparable to that observed for ciTotiSCs and TLSCs (**Fig. 1g**). Consistent with this observation, 36 h-induced cells showed increased expression of a set of genes related to maternal transcripts, zygotic genome activation (ZGA), and totipotency compared with parental mESCs and several previously reported 2C-like cell populations (**Fig. 1h**).

Collectively, these findings indicate that our 2C induction system robustly drives the mESCs toward a molecular state that shares key transcriptional and epigenomic features with 2C-stage embryos. The high efficiency of the system therefore provides a useful platform for investigating the molecular mechanisms underpinning the reprogramming from pluripotency to a totipotent-like state.

### scRNA-seq characterization of branched trajectory in chemical reprogramming

To gain deeper insights into the cellular heterogeneity and molecular features of CIM-treated cells, we analyzed time-series scRNA-seq data obtained at 0 h, 12 h, 24 h, and 36 h of CIM treatment. We pooled all cells and conducted clustering analysis, which identified ten major cell clusters (**Fig. 2a**). Clusters 0h_C1, 0h_C2 and 0h_C3 represent mESCs as they expressed pluripotency genes such as *Pou5f1*, *Nanog*, *Sox2*, *Klf4*, *Esrrb*, and *Tfcp2l1* (**Fig. 2a; Supplementary Fig. S3a, b**). Cluster 12h_C1, 12h_C2, and 12h_C3 appear to be an intermediate cell population, showing reduced expression of pluripotency genes (e.g. *Klf4* and *Nanog*) with detectable but low expression of 2-cell-embryo-specific transcripts such as *Zscan4* genes (**Fig. 2a; Supplementary Fig. S3a, b**). The 24 h and 36 h induced cells were primarily divided into two major clusters respectively, with each exhibiting distinct marker gene expression patterns (**Fig. 2a; Supplementary Fig. S3a, b**). Clusters 24h_C2 and 36h_C2 appeared to be totipotent-like cells with high expression of *Zscan4*, *Usp17lc*, and *Tcstv3*, whereas these markers were expressed at much lower levels in clusters 24h_C1 and 36h_C1 (**Fig. 2b; Supplementary Fig. S3a, b**). Concordantly, Pearson correlation analysis revealed that clusters 24h_C2 and 36h_C2 were similar to 2-cell embryos, whereas clusters 24h_C1 and 36h_C1 displayed moderate similarities to various developmental stages of mouse embryos from 2-cell to blastocyst (**Fig. 2c**). As expected, trajectory analysis identified a sequential branching during chemical reprogramming, ultimately giving rise to one reprogramming potential fate (36 h tdT-high) and one non-reprogramming fate (36 h tdT-low) (**Fig. 2d; Supplementary Fig. S3c-f**). These findings suggest that chemical induction leads to branching into multiple trajectories, representing distinct reprogrammed and non-reprogrammed cell fates.

**Fig. 2.**
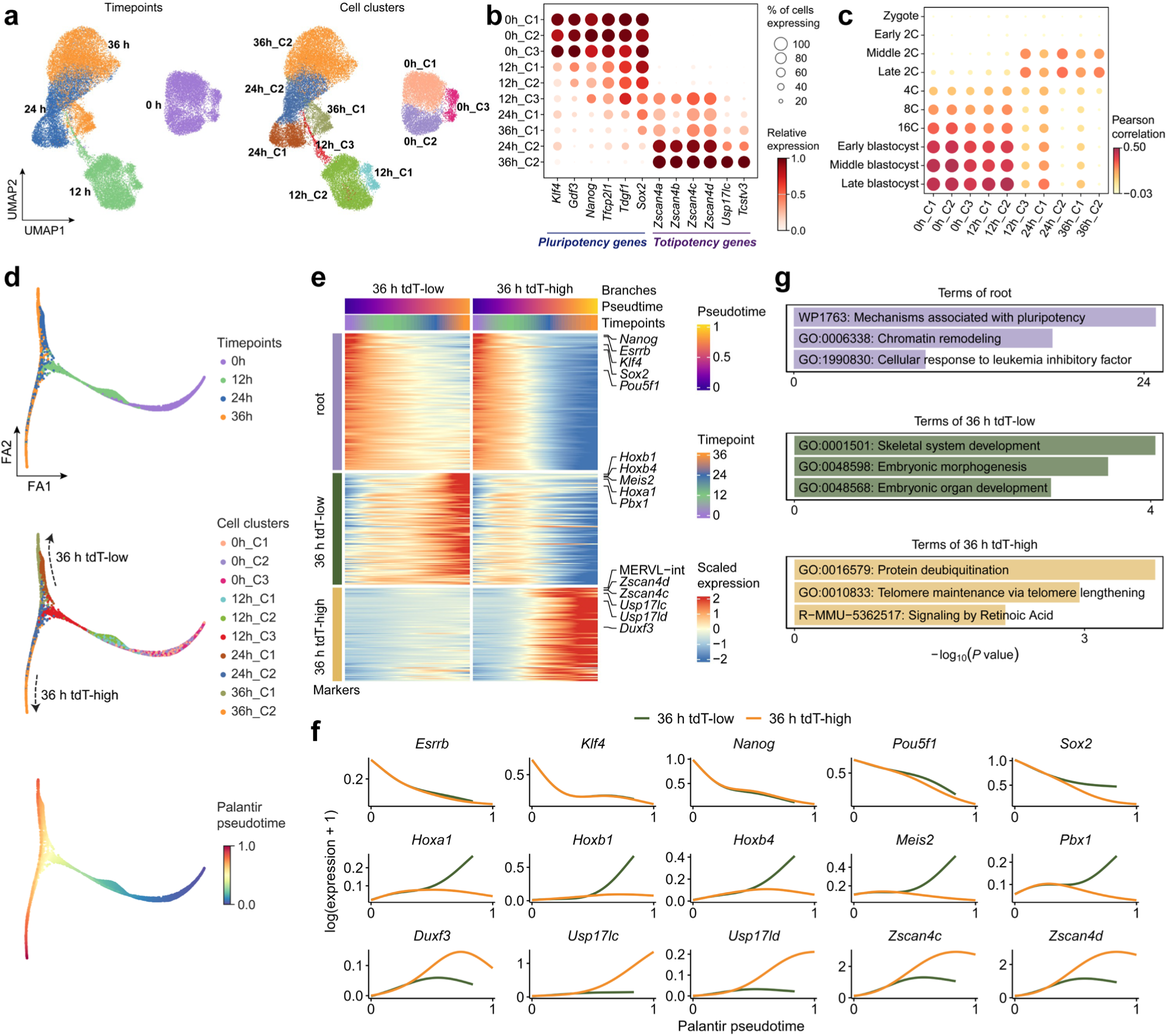
scRNA-seq identified different branching trajectories of cells during 2C-like reprogramming. **a**, Uniform manifold approximation and projection (UMAP) analysis of integrated scRNA-seq profiles across four chemical reprogramming timepoints, with cells colored by timepoint on the left and by the Leiden clustering on the right. **b**, Expression of pluripotency and totipotency genes across the ten cell clusters shown in (**a**). Dot size indicates the percentage of cells in each cluster expressing the gene, and color intensity reflects the scaled expression values. **c**, Pearson correlation analysis of pseudo-bulk RNA-seq data from ten chemical reprogramming cell clusters and various stages of early mouse embryos (from zygote to late blastocyst), based on curated totipotency and pluripotency gene sets. **d**, ForceAtlas2 layout visualizing the different reprogramming trajectories, with cells colored by chemical reprogramming timepoints (upper), cell clusters (middle), and Palantir pseudotime (bottom). The upper and lower trajectories represent the 36 h tdT-low and 36 h tdT-high branches, respectively. **e**, Heatmap showing the dynamics of gene expression along Palantir pseudotime for the 36 h tdT-low, and 36 h tdT-high trajectories. Each row represents a differentially expressed gene or transposable element (TE), collectively referred to as markers, identified between root, 36 h tdT-low and 36 h tdT-high cell groups, as defined in **Supplementary Fig. S3g**. **f**, Comparison of gene expression trends along Palantir pseudotime for genes from the enriched pathways in root, 36 h tdT-low, and 36 h tdT-high cell groups. **g**, Bar plots showing pathway enrichment of marker genes for the root, 36 h tdT-low, and 36 h tdT-high cell groups.

Next, we delineated the molecular signatures distinguishing reprogramming from non-reprogramming outcomes. We compared the gene expression profiles across the three defined cell states: the root cells, and the two terminal populations (36 h tdT-low, and 36 h tdT-high) (Methods; **Supplementary Fig. S3g**). We observed that pathways such as skeletal system development and embryonic organ development were significantly enriched in 36 h tdT-low branch, with *Hoxa1*, *Hoxb1*, *Hoxb4*, *Meis2*, and *Pbx1* expressed at high levels (**Fig. 2e-g**). This suggests that these cells may have already entered a differentiation program. By contrast, the 36 h tdT-high branch was enriched for the protein deubiquitination pathway and displayed elevated expression of not only totipotency genes *Duxf3* and *Zscan4c/d*, but also *Usp17l* family members (**Fig. 2e-g**), whose overexpression has been reported to reduce H2AK119ub1 levels and promote activation of *Duxf3* and 2C genes^26^. In addition, analysis of the expression of genes encoding chromatin architectural proteins and other chromatin regulators along Palantir pseudotime across the two branches revealed a more pronounced down-regulation in the 36 h tdT-high branch, consistent with a relaxed 3D genome during the pluripotency-to-totipotency transition^13, 27^ (**Supplementary Fig. S3h**).

### SP2 impedes the pluripotent-to-totipotent state transition

A fraction of cells failing to complete the 2C-like transition suggests the presence of factors promoting differentiation, thus hindering the pluripotency-to-totipotency transition. To identify potential barriers, we inferred potential upstream transcription factors based on the marker genes of the ten distinct cell clusters (**Fig. 3a**). Notably, motifs associated with the specificity protein (SP) family were highly enriched in cluster 36h_C1, and the 24 predicted target genes of SPs were significantly enriched in biological processes associated with rhombomere development, embryonic morphogenesis, and anterior/posterior pattern specification, implying that these factors may impede reprogramming by promoting alternative developmental programs (**Fig. 3a; Supplementary Fig. S4a**).

**Fig. 3.**
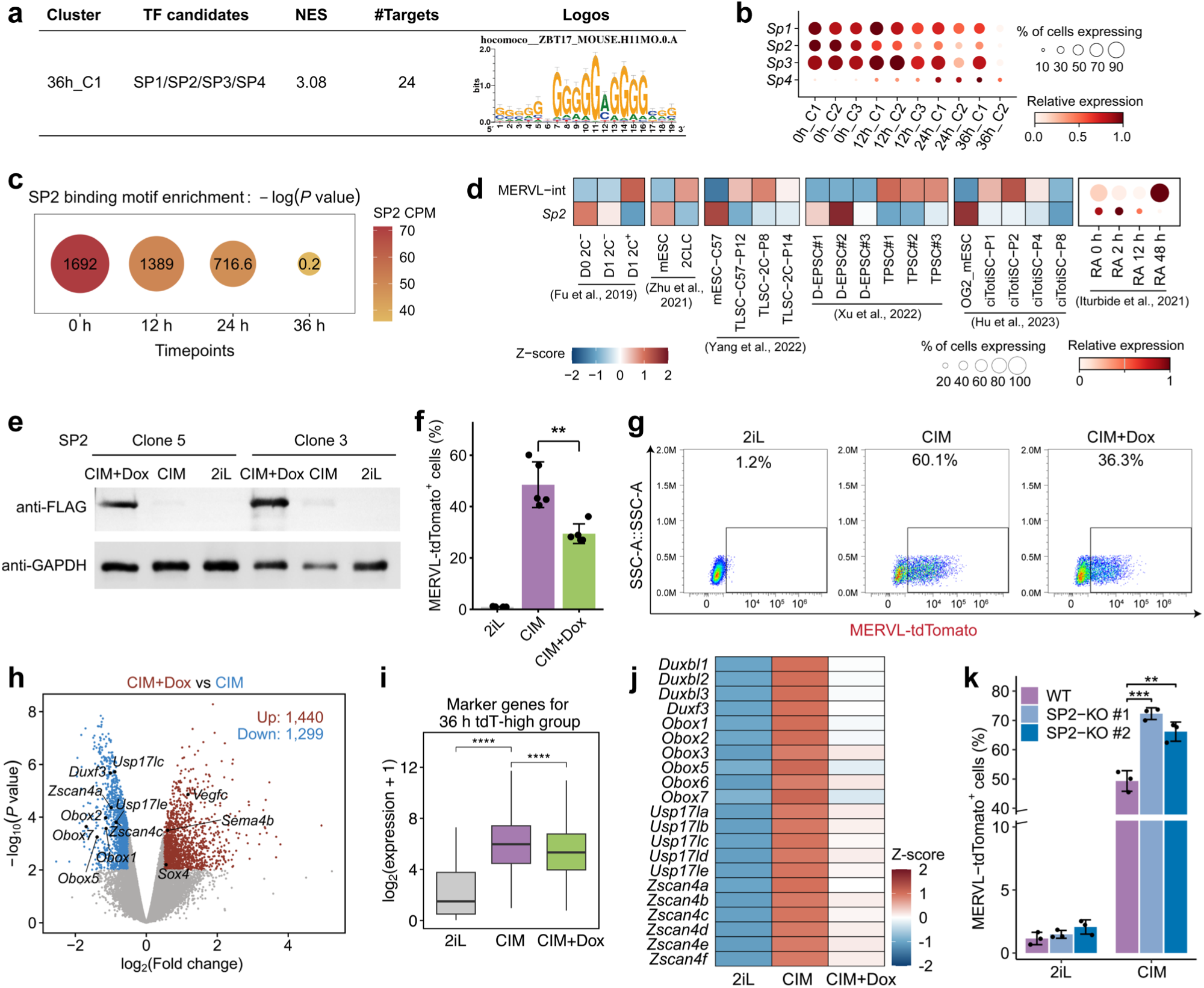
SP2 acts as a barrier to 2C-like reprogramming. **a**, Potential TF motif enrichment in the 36h_C1 cell cluster. NES, normalized enrichment score. **b**, Dot plot showing the expression patterns of *Sp* family genes in different cell clusters identified from scRNA-seq. The dot size represents the percentage of cells in each cluster expressing the corresponding genes, and the color intensity reflects the scaled expression values. **c**, SP2 motif enrichment in ATAC-seq peaks at 0 h, 12 h, 24 h and 36 h after 2C-like induction. Dot size represents -log(*P* value). Dot color indicates *Sp2* expression levels (CPM). **d**, Heatmap showing the comparative expression of MERVL-int and Sp2 across different cell populations. D0 2C^−^, mESCs; D1 2C^−^, intermediate-state cells; D1 2C^+^, *Dux*-induced 2CLCs; RA, retinoic acid. RA 0, 2, 12 and 48 h indicate mESCs grown in LIF and treated with RA for 0, 2, 12 and 48 h, respectively. **e**, Western blot showing SP2 protein levels after 36 h of culture in 2iL medium and in CIM with or without Dox. **f**, The percentage of MERVL-tdTomato^+^ cells after 36 h of culture in 2iL medium or CIM with or without Dox, quantified by FACS in four independent mESC clones. Shown are mean ± s.d., *n* = 5 biologically independent cell cultures. \*\**P* < 0.01; two-tailed unpaired Student’s *t*-test. **g**, Representative scatter plot for the experiment in (**f**), showing MERVL-tdTomato fluorescence measurements of individual cells as assayed by FACS. **h**, Volcano plot of differentially expressed genes following SP2 overexpression in CIM for 36 h. Red points: upregulated (log_2_FC ≥ 0.5, *P* < 0.01); Blue points: downregulated (log_2_FC ≤ -0.5, *P* < 0.01). **i**, Box plots showing log2 expression levels of marker genes specific to the 36 h tdT-high cell group across cells cultured in 2iL medium or in CIM with or without Dox. \*\*\*\**P* < 0.0001; two-tailed Wilcoxon signed-rank test. **j**, Heatmap showing relative expression levels of selected totipotency genes across 2iL, CIM, and CIM + Dox conditions. **k**, Percentage of MERVL-tdTomato^+^ cells in wild-type (WT) and SP2 KO mESCs cultured in 2iL medium and CIM. Shown are mean ± s.d., *n* = 3 independent experiments. \*\**P* < 0.01, \*\*\**P* < 0.001; one-tailed unpaired Student’s *t*-test.

We next examined the expression patterns of individual SP family members during 2C-like induction. Among these factors, *Sp2* showed relatively low expression in cluster 36h_C2, consistent with its low expression across the 36 h tdT-high branch. (**Fig. 3b; Supplementary Fig. S4b-d**). Consistent with the transcription changes, ATAC-seq revealed higher enrichment of the SP2 motif in accessible regions at 0 h than at 36 h, paralleling the decrease in *Sp2* expression during induction (**Fig. 3c; Supplementary Fig. S4c, d**). Furthermore, we examined *Sp2* expression across previously reported totipotent-like cell models. Compared with mESCs, *Sp2* expression was consistently reduced in several independently established totipotent-like populations including *Dux*-induced 2CLCs^9^, spontaneously converted 2CLCs^27^, TLSCs^24^, TPSCs^17^, ciTotiSCs^16^, and RA-induced 2CLCs^28^ (**Fig. 3d**). Thus, down-regulation of *Sp2* is not restricted to our chemical induction system but also observed across different approaches for inducing a totipotent-like state. Based on these observations, we hypothesized that SP2 may act as a barrier to the pluripotency-to-totipotency transition.

To test this hypothesis, we generated doxycycline (Dox)-inducible SP2-overexpressing mESC lines, and in parallel, established inducible SP1- and SP3-overexpressing mESC lines for the comparison. Upon induction with our system, Dox-induced SP2 overexpression significantly reduced the proportion of MERVL-tdTomato^+^ cells across four independent mESC clones (**Fig. 3e-g**). However, overexpression of SP1 or SP3 had a subtle effect on the percentage of MERVL-tdTomato^+^ cells under the same conditions, with SP3 slightly increasing 2C-like reprogramming efficiency (**Supplementary Fig. S4e, f**). Consistent with these findings, SP2 overexpression significantly downregulated marker genes of the 36 h tdT-high cell group, as well as key 2C-associated genes, including *Duxf3*, *Duxbl*, *Obox*, *Usp17l*, and *Zscan4* families (**Fig. 3h-j**). These results indicate that the inhibitory effect observed is more pronounced for SP2 but not other SP members. We next generated two independent *Sp2*-KO mESC lines using CRISPR-Cas9 (**Supplementary Fig. S4g**) and subjected them to 2C-like induction. As expected, loss of SP2 further increased the proportion of MERVL-tdTomato^+^ cells compared with control mESCs (**Fig. 3k; Supplementary Fig. S4h**). Together with the gain-of-function results, these findings demonstrate that SP2 functions as a barrier to the pluripotent-to-totipotent transition in embryonic stem cells.

### SP2 impacts 2C-like induction and promotes differentiation by activating *HOX* gene family in an RA-dependent manner

To investigate how SP2 impedes the acquisition of the 2C-like state, we compared the transcriptomes of control and SP2-deficient mESCs during 2C-like induction. SP2 deletion resulted in approximately 2,000 differentially expressed genes (**Fig. 4a**). Upregulated genes were significantly enriched in marker genes of the 36 h tdT-high cell group, whereas downregulated genes were significantly enriched in marker genes of the 36 h tdT-low cell group (*P* < 0.05, hypergeometric test for overrepresentation) (**Fig 4a; Supplementary Fig S5a**). Notably, the downregulated genes were enriched for multiple developmental processes, including skeletal system development, anterior/posterior pattern specification, and embryonic organ morphogenesis (**Fig. 4b**). Moreover, GSEA showed that SP2 deletion attenuated pattern specification-related developmental programs (**Supplementary Fig. S5b**). In particular, several developmental genes, including members of the *Hox* family that were preferentially expressed in 36 h tdT-low cells, were downregulated upon SP2 deletion (**Fig. 4a**). In parallel, genes associated with the 2C-like state, including *Obox* family genes, *Duxf3*, and *Zscan4a*, were upregulated following SP2 deletion (**Fig. 4a**). These transcriptional changes were consistent with the increased frequency of MERVL-tdTomato^+^ cells observed in SP2-deficient mESCs, suggesting that SP2 may initiate developmental gene activation during 2C-like reprogramming.

**Fig. 4.**
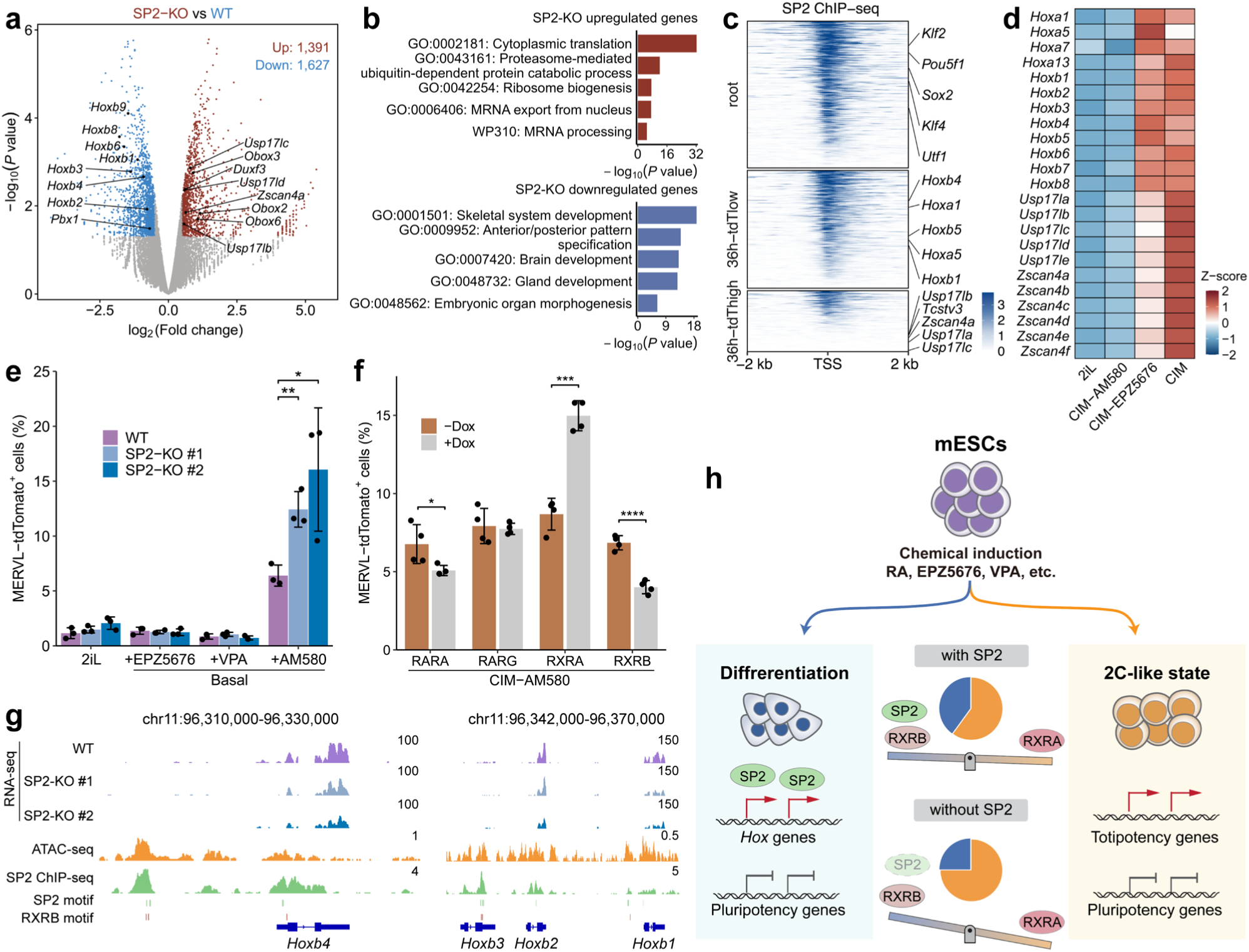
SP2 impacts 2C-like induction and promotes differentiation by activating *Hox* gene family in an RA-dependent manner. **a**, Volcano plot showing differentially expressed genes in SP2-KO versus WT mESCs cultured in CIM for 36 h. Red points: upregulated (log_2_FC ≥ 0.5, *P* < 0.05); Blue points: downregulated (log_2_FC ≤ -0.5, *P* < 0.05). Selected *Hox* family members and 2C-associated genes are highlighted. **b**, Metascape bar plots showing enriched pathways among the upregulated and downregulated genes identified in (**a**). **c**, Heatmap showing SP2 ChIP-seq signal at the promoter regions of marker genes in the root, 36 h tdT-low, and 36 h tdT-high cell groups. **d**, Heatmap showing the relative expression of *Hox*, *Usp17l*, and *Zscan4* family genes in mESCs cultured in 2iL medium, CIM without AM580, CIM without EPZ5676, and CIM. **e**, Percentage of MERVL-tdTomato^+^ cells in WT and two SP2-KO mESCs cultured in 2iL medium or basal medium supplemented with EPZ5676, VPA, or AM580. Shown are mean ± s.d., *n* = 3 independent experiments. \**P* < 0.05, \*\**P* < 0.01; one-tailed unpaired Student’s *t*-test. **f**, Percentage of MERVL-tdTomato^+^ cells of RARA, RARG, RXRA, and RXRB overexpressing mESCs in CIM lacking AM580 with or without Dox. Shown are mean ± s.d., *n* = 4 independent experiments. \**P* < 0.05, \*\*\**P* < 0.001; \*\*\*\**P* < 0.0001; two-tailed unpaired Student’s *t*-test. **g**, IGV screenshot showing SP2 ChIP-seq, RNA-seq, ATAC-seq, and motifs enrichment at representative *Hox* genes. **h**, Graphic summary of the biological functional role of SP2 during 2C-like induction in the presence of RA signaling.

SP2 was known as a transcription factor that directly binds chromatin and regulates gene expression. We next investigated the genome-wide occupancy of SP2 in CIM-induced cells. ChIP-seq revealed that SP2 predominantly binds to promoter regions (transcription start site ± 2 kb), with strong enrichment of the canonical SP2 motif (**Supplementary Fig S5c, d**). Of note, SP2 occupied a large proportion of promoters of marker genes specific to the root and 36 h tdT-low populations (∼70-80%), whereas binding was less frequent at promoters of marker genes specific to the 36 h tdT-high population (∼40%) (**Fig. 4c; Supplementary Fig S5e**). Consistent with the transcriptional changes observed following SP2 deletion, SP2 binding was detected at multiple developmental gene loci, including several *Hox* genes (**Fig. 4c**), supporting a direct role for SP2 in regulating the developmental program that competes with 2C-like induction.

Previous studies have reported that RA treatment can induce a 2C-like state in mESCs while simultaneously promoting differentiation-associated gene expression^28^. Consistent with these observations, omission of AM580 from our 2C-like induction medium markedly reduced the expression of *Hox* family genes as well as 2C-associated genes (**Fig. 4d; Supplementary Fig S5f-h**). In contrast, removal of EPZ5676 caused little effect on *Hox* gene expression (**Fig. 4d; Supplementary Fig S5f-h**), suggesting that the developmental program associated with *Hox* gene activation is closely linked to RA signaling during 2C-like induction. We therefore tested whether the effect of SP2 on 2C-like induction depended on RA receptor activation. Control and SP2-KO mESCs were cultured in basal medium supplemented with AM580, EPZ5676, or VPA individually. Strikingly, SP2 deletion significantly increased the proportion of MERVL-tdTomato^+^ cells following AM580 treatment, whereas no clear increase was observed following EPZ5676 or VPA treatment (**Fig. 4e; Supplementary Fig. S4h**). RA signaling is mediated by six receptors that are classified into two subfamilies, RARs and RXRs. We first examined the expression levels of individual RA receptors during 2C-like induction, and found that only *Rara*, *Rarg*, *Rxra*, and *Rxrb* were expressed (**Supplementary Fig S5i**). Notably, *Rxra* expression was increased in the 36h tdT-high branch, whereas *Rara*, *Rarg*, and *Rxrb* were more enriched in 36h tdT-low branch, suggesting distinct roles of RA receptors in 2C-like induction (**Supplementary Fig S5j**). To investigate how different RA receptors affect 2C-like reprogramming, we individually overexpressed RARA, RARG, RXRA, or RXRB during 2C-like induction in the absence of AM580. RARA and RARG had subtle effects on 2C-like induction, whereas RXRA overexpression increased the proportion of MERVL-tdTomato^+^ cells and RXRB overexpression inhibited 2C-like induction (**Fig. 4f; Supplementary Fig S5k**). We next tested whether RXRB might be functionally linked to SP2 at developmental loci. At representative SP2-bound *Hox* gene loci, RXRB motifs were found in close proximity to SP2 motifs (**Fig. 4g**). These findings raise the possibility that RXRB and SP2 cooperate at *Hox* regulatory regions to promote the developmental program that competes with 2C-like induction. Altogether, these findings reveal that the transition toward the 2C-like state is shaped by a balance between totipotency-associated and developmental programs, with SP2 acting at the intersection of RA signaling and developmental gene activation (**Fig. 4h**).

## Discussion

The transition from pluripotency to totipotency is accompanied by extensive changes in transcriptional programs and cell fate potential. Although several approaches have been developed to induce 2C-like cells from mESCs, the factors that determine whether cells successfully acquire totipotent-like features or instead enter alternative developmental states remain poorly understood. Our study identifies SP2 as a regulator of this cell fate choice in an RA-dependent manner. This finding provides a possible explanation for the diverse responses of pluripotent cells to RA activation and suggests that acquisition of totipotent-like features is influenced not only by activation of a totipotency program, but also by suppression of competing developmental programs.

RA signaling is well established as a major regulator of embryonic patterning and lineage specification, yet its role in totipotency induction presents an apparent paradox. RA receptor activation can promote the emergence of 2C-like cells, while at the same time activating developmental genes, including *Hox* genes, that are characteristic of later embryonic patterning. Our findings suggest that these activities may represent competing outputs of the same signaling pathway, and SP2 provides a potential molecular link between these distinct outputs. SP2 is a member of the specificity protein (SP) family of transcription factors, characterized by three highly conserved C2H2-type zinc fingers that recognize GC-rich DNA motifs. Previous reports have demonstrated important roles for SP2 in both neurogenesis and tumorigenesis^29–32^. However, compared with other SP family members, the functional contribution of SP2 during early embryonic development remains poorly understood. We propose that, upon RA receptor activation, SP2 contributes to the activation of developmental transcriptional programs, thereby favoring differentiation over acquisition of the 2C-like state. The preferential effect of SP2 loss on AM580-induced reprogramming further suggests that SP2 does not act as a general inhibitor of totipotency, but may instead function as a context-dependent effector of RA signaling. In addition, SP2 is unlikely to be the sole determinant of the differentiation branch, as SP2 overexpression only partially reduced 2C-like induction. Other transcription factors and signaling pathways may therefore cooperate with SP2 to determine the balance between totipotency and differentiation. Nevertheless, our findings reveal a previously unrecognized connection between SP2 and RA signaling and suggest that developmental transcription factors can act as gatekeepers that determine whether pluripotent cells respond to a developmental signal by acquiring broader potency or by committing to differentiation.

## Materials and Methods

### Cell culture

The mESCs containing MERVL-LTR-tdTomato reporter was a gift from Wang’s lab. All the mESCs were cultured in mESC medium (DMEM High-glucose (HyClone, SH30022.01) supplemented with 15% FBS (Lonsera, S711-001s), NEAA (Gibco,11140076), Glutamax (Gibco, 35050079), Sodium Pyruvate (Gibco, 11360070), 2-Mercaptoethanol (Sigma, M3148-100ML), Leukemia Inhibitory Factor (novoprotein, C690), MEK inhibitor PD0325901 (1 µM, DC CHEMICALS, DC1056), and GSK3 inhibitor CHIR99021 (3 µM, DC CHEMICALS, DC1023)) on 0.1% gelatin coated plates. For 2C-like cell induction, mESCs were switched to chemical induction medium (CIM) containing DMEM High-glucose, FBS (Gibco, 10099141), KnockOut SR (Gibco, 10828028), B27 (Gibco, 17504044), NEAA, 2-Mercaptoethanol (referred to as the basal medium), supplemented with 100 nM AM580 (Selleck, S2933), 1 μM BML277 (TargetMOL, T2033), 1 μM JNK inhibitor VIII (MCE, HY-107598-1mg), 100 μM VPA (TargetMol, T7064), 5 μM EPZ5676 (Selleck, S7062), 5 mM LiCl (Sigma, 793620-500G), and 50 μg/mL Vitamin C (Sigma, 49752) for 36 h. For single addition, each small molecule was added into basal medium for 2C-like induction.

For establishing overexpression cell lines, PB transposon plasmid containing TRE-controlled SP1, SP2, and SP3, together with transposase, were transfected into 2C-reporter mESCs. For generating SP2 knockout cell lines, pX330 targeting SP2 exon2 were transfected into 2C-reporter mESCs. Single colonies were hand-picked for genotyping, and the mutations were detected with sanger-sequencing.

### FACS analysis

Cells were trypsinized and harvested into tubes, and then washed two times with cold PBS before flow cytometry analysis. Cells were loaded and analyzed by Beckman Coulter CytoFLEX S and the collected data were analyzed by the software FlowJo (version 10.8.1).

### Western blot

Cells were lysed in cell lysis buffer (50 mM Tris pH7.5, 150 mM NaCl, 1% NP-40, 1mM EDTA, 0.1% SDS, and Protease inhibitor cocktail (Roche, 05056489001)) on ice. Total cell extracts were separated on a 10% SDS-Page (Yamei, PG112), and transferred to a PVDF Membrane (Millipore, IPVH00010). After blocking with 5% non-fat milk, the following primary antibodies were used: Flag (1:1000, Sigma, F1804), SP2 (1:1000, proteintech, 25000-1-AP) and Gapdh (1:5000, Aksomics, KC-5G5). Goat anti-mouse HRP (1:5000, Aksomics, KC-MM-035) was used as secondary antibody. Membranes were analyzed on MINICHEMI chemiluminescent imaging system.

### Real-time Quantitative PCR and RNA-seq library preparation

Total RNAs were extracted from cells with TRIzol following the standard protocol. 1 µg of total RNA per sample was converted into cDNAs using HiScript III RT SuperMix for qPCR (+gDNA wiper) kit (Vazyme, R323), and then analyzed by qPCR with Cham QTM qPCR SYBR® Green Master Mix (Vazyme, Q311-03) on Bio Rad cfx connect real-time system. For polyA-based RNA-seq, VAHTS mRNA Capture Beads (Vazyme, N401-2) and VAHTS Universal V8 RNA-seq Library Prep Kit for Illumina (Vazyme, NR605) were used for RNA library construction, and the libraries were sequenced at HUAYIN HEALTH. All the primers used for qPCR can be found in **Supplementary Table S1**.

### ATAC-seq library preparation

A total number of 50,000 cells per sample were used for ATAC-seq. ATAC-seq libraries were prepared using Hyperactive ATAC-Seq Library Prep Kit for Illumina (Vazyme, TD711-01) following the standard protocol. ATAC-seq libraries were sequenced at HUAYIN HEALTH.

### Single-cell RNA-seq library preparation

Single-cell suspensions were loaded onto 10× Genomics Chromium v3.1 system to generate gel beads-in-emulsion (GEMs), where all full-length cDNA molecules within a GEM shared a common 10× barcode. Following incubation, GEMs were disrupted, and cDNA was PCR-amplified. For library construction, 10 μL (25% of total cDNA) was used to generate single-cell 3’ gene expression libraries, which were then purified using SPRIselect beads. Library quality was assessed by analyzing size distribution (Qsep100) and quantifying concentration (Qubit 4.0 fluorometer). Finally, sequencing was performed on BGI T7 system.

### ChIP-seq library preparation

Cells were cross linked with 1% formaldehyde (Sigma, 252549) for 10 min at room temperature and glycine (125 mM) was used to quench cross-linking for 5 min. Cells were then washed twice with cold PBS and lysed with ChIP Buffer (50 mM Tris–HCl (pH 7.9), 500 mM NaCl, 0.1% DOC, 0.1% SDS, 2 mM EDTA, 1% Triton X-100, Protease inhibitor cocktail (Roche, 05056489001)). Then, the chromatin was sheared by sonication (Qsonica) under the following conditions, 30 s/30 s on/off, 100% amplitude, 2 min per cycle, 4-5 cycles. Sheared chromatin was first incubated with 2μg Flag Antibody (Sigma, F1804) at 4 °C for at least 6 h, and then Protein A and G Beads (Life Technologies; 10002D, 10003D) were added and incubated with chromatin-antibody mix at 4 °C overnight. After immunoprecipitation, the mixture was washed with buffer 1(50 mM Tris-HCl (pH 7.9), 500 mM NaCl, 1% Triton X-100, 0.1% DOC, 1 mM EDTA), buffer 2(10 mM Tris-HCL (pH 7.9), 250 mM LiCl, 0.5% NP40, 0.5%

DOC, 1 mM EDTA), and buffer3 (50 mM Tris-HCL (pH 7.9)). Samples were de-crosslinked with 1% SDS at 65 °C, and then treated with RNase A and proteinase K at 56 °C. DNA was extracted using mini elute Kit (Qiagen) and used for further analysis. Universal DNA Library Prep Kit for Illumina (Vazyme, V3ND607-02) was applied for ChIP library construction, and the ChIP libraries were sequenced at HUAYIN HEALTH.

### Public datasets

The public datasets used in this study, including ATAC-seq, RNA-seq, and scRNA-seq, are summarized in **Supplementary Table S2**.

### RNA-seq and Smart-seq/Smart-seq2 scRNA-seq data preprocessing

Raw bulk RNA-seq data and public preimplantation embryo scRNA-seq data (GSE45719) generated using Smart-seq/Smart-seq2 were preprocessed as follows. The raw reads were subjected to trim_galore (version 0.6.10, https://www.bioinformatics.babraham.ac.uk/projects/trim_galore/) to remove adaptors and low-quality reads. The clean reads were aligned to the mm10 mouse genome with mouse gene annotation based on GENCODE vM21 using STAR (version 2.7.10b)^33^, with parameters “--outFilterMultimapNmax 100 --winAnchorMultimapNmax 100 --outMultimapperOrder Random --runRNGseed 777 --outSAMmultNmax 1 --readFilesCommand zcat”. Transcript and TE expression levels were quantified using TEcount command in TEtranscripts package (version 2.2.3)^34^.

### Differential expression analysis of bulk RNA-seq data

For bulk RNA-seq samples, gene and repeat count matrices were normalized using the “Trimmed Mean of M values” (TMM) method^35^ in edgeR package (version 3.32.1)^36^ and transformed to counts per million (CPM) using the cpm function. Differential expression analysis was assessed with edgeR, employing the generalized linear model-based approach for samples with replicates and the exactTest function for those without replicates.

### Pearson correlation analysis of RNA-seq data

Pairwise Pearson correlations between RNA-seq samples were calculated using the personr function from the scipy.stats package based on normalized CPM expression values of genes from curated totipotency and pluripotency gene sets^24^.

### Gene set enrichment analysis (GSEA)

Gene Set Enrichment Analysis (GSEA) was performed using fgsea R package^37^. For the comparison of CIM-treated cells at 36 h versus 0 h, the 2 cell and pluripotency gene sets^23^, and the totipotency gene set^24^ were analyzed. For the SP2-KO versus WT comparison, gene sets from the M2 and M5 collections of Mouse MSigDB (v2026.1.Mm) were analyzed. *P* values were adjusted using the Benjamini-Hochberg method. *P* values and Benjamini-Hochberg-adjusted *P* values (FDRs) were calculated.

### 10× Genomics single-cell RNA-seq analysis

#### Processing of sequencing files and cell filtering

The reads were aligned to the mm10 genome with mouse gene annotation based on GENCODE vM21 using STARsolo function of STAR, with the parameters “--outSAMattributes NH HI AS nM CR CY UR UY --outFilterMultimapNmax 100 --winAnchorMultimapNmax 100 --outMultimapperOrder Random --runRNGseed 777 --outSAMmultNmax 1 -- readFilesCommand zcat”. The scTE (version 1.0)^38^ processing pipeline was used to generate per-cell molecule count matrices for genes and transposable elements based on gene annotations (GENCODE vM21) and transposable elements (rmsk mm10). Cells with fewer than 10 UMIs or fewer than 10 detected genes were filtered out, and the top 20,000 cells with the highest gene count were retained.

Downstream analysis was performed using the Scanpy package (version 1.10.3), with low-quality cells filtered based on mitochondrial gene percentage, total counts, and the number of detected genes. Scrublet (scanpy, external.pp.scrublet, default parameters)^39^ was used to predict and remove potential doublet cells. After quality control of the scRNA-seq data from CIM-treated cells, approximately 9,000 cells per sample were retained for downstream analysis. The 0 h, 12 h, 24 h and 36 h datasets were concatenated using Scanpy into a single AnnData object for subsequent analysis.

#### Cell clustering and visualization

Total counts per cell were normalized to 10,000, and then log1p was applied. Highly variable genes (HVGs) were identified using thresholds of min_mean = 0.05, max_mean = 4, and min_disp = 0.6. PCA was performed on the normalized matrix of HVGs (scanpy, tl.pca, svd_solver = “arpack”, n_comps = 50, use_highly_variable=True). The K nearest-neighbours (KNN) graph (scanpy, pp.neighbors, n_neighbors = 25, n_pcs = 50) was generated from the PCA space, and was used as basis for cluster identification using Leiden algorithm (scanpy, tl.leiden, res = 0.5), as well as UMAP embedding (scanpy, tl.umap, default parameters). Cell clusters were labeled according to the predominant sampling timepoint and cluster number, denoted as timepoint_C# (e.g., 0h_C1).

### Transcription factor inference from cell cluster-specific marker genes

To identify cell cluster-specific differentially expressed genes, each cluster was randomly downsampled to 500 cells without replacement to mitigate cluster-size bias. Marker discovery was then performed in Scanpy using tl.rank_genes_groups with the Wilcoxon rank-sum test, requesting the top 400 genes per cluster against the remaining cells. Differentially expressed genes were retained if *P* < 0.05, log2 fold change > 0.5, detection fraction in the target cluster > 0.2, and detection-fraction difference (target - rest) > 0.15. Potential key TFs regulating cell cluster-specific gene networks were inferred using the RcisTarget package (version 1.20.0)^40^. Specifically, the marker genes from each cell cluster were analyzed for enriched TF binding motifs within a 20 kb window centered on the transcription start site (TSS). Motifs with Normalized Enrichment Score (NES) > 3 were considered significant. TFs associated with these enriched motifs were considered potential key regulators only if they were expressed in at least 10% of the cells within the respective group. A representative enriched motif was displayed in **Fig. 3a**, together with its corresponding cell cluster, associated TF candidates, NES, number of target genes, and sequence logo.

### Trajectory inference

To balance the cell numbers of different clusters, each cluster was randomly downsampled using random.choice function in python to 500 cells without replacement. Top 1,000 highly variable genes were used to perform PCA (scanpy, tl.pca, svd_solver = “arpack”, n_comps = 50, use_highly_variable = True). Trajectory inference was performed by Harmony (version 0.1.5)^41^ and Palantir (version 1.4.1)^42^ as described previously^43^. Briefly, the first 50 principal components and CIM induction time annotation were used for aligning the data using Harmony package. Augmented affinity matrix was generated (harmony, core.augmented_affinity_matrix, n_neighbors = 50). Diffusion map was then generated from this affinity matrix (palantir, utils.run_diffusion_maps, n_components = 10) and multiscale space was determined (palantir, utils.determine_multiscale_space, n_eigs = 3). To generate the Force Atlas embedding, a t-SNE embedding was first generated from the multiscale diffusion space (scanpy, tl.tsne, perplexity = 40, learning_rate = 1000). Second, from the same multiscale diffusion space, a KNN graph was generated (scanpy, pp.neighbors, n_neighbors = 50). Force atlas was then generated from the neighbors graph using the tSNE coordinates as initialization (scanpy, tl.draw_graph, init_pos = “X_tsne”, maxiter = 3000).

For pseudotime trajectory analysis, the cell with the highest FA1 coordinate in cluster 0h_C1 was selected as the root cell. The cells with the highest and lowest FA2 coordinates in clusters 36h_C1 and 36h_C2, respectively, were selected as the two terminal states (**Supplementary Fig. S3c**). Pseudotime and branch probabilities to each terminal state were inferred by palantir.core.run_palantir (num_waypoints = 500) using the root cell and terminal states. For heatmap visualization, gene expression values were smoothed along Palantir pseudotime using the R package gam (version 1.22.5), and heatmaps of the smoothed expression trends for each trajectory were generated with ComplexHeatmap (version 2.16.0). MAGIC (Markov Affinity-based Graph Imputation of Cells) imputation was conducted to denoise the gene expression matrix (palantir, utils.run_magic_imputation, default parameters). Gene expression trends for selected genes were computed along Palantir pseudotime for each branch using the MAGIC-imputed gene expression matrix (palantir, presults.compute_gene_trends, expression_key = “MAGIC_imputed_data”).

### Differential expression analysis in pseudotime-defined cell groups

To enable downstream comparative analyses of gene expression and functional enrichment, we partitioned the reprogramming trajectories into three representative cell groups: root cells, defined as the 500 cells with the lowest pseudotime values, and 36 h tdT-low and 36 h tdT-high terminal cells, defined as the 500 cells with the highest pseudotime values along each respective branch (**Supplementary Fig. S3g**). Differential expression analysis was performed in Scanpy using tl.rank_genes_groups with the Wilcoxon rank-sum test, retrieving the top 800 genes per group. Genes or TEs were defined as group-specific markers if log2 fold change > 1, detection fraction in the target cluster > 0.3, detection-fraction difference (target - rest) > 0.1, and *P* < 0.05. For downstream analyses, markers identified in more than one group were assigned to the group in which they showed the highest log2 fold change.

### Gene ontology term enrichment analysis

Gene Ontology enrichment analysis was performed using Metascape (http://metascape.org), which utilizes the hypergeometric test and Benjamini-Hochberg *P*-value adjustment^44^. The list of genes was uploaded to Metascape as a text file. The “Express Analysis” option was selected after setting both the “Input as species” and “Analysis as species” parameters to M. musculus.

### ATAC-seq analysis

#### Data processing and peak calling

Raw reads were processed with trim_galore (version 0.6.10) to remove adaptors and low-quality reads. The trimmed reads were mapped to the mm10 reference genome using bowtie2 (version 2.5.4)^45^, with parameters “--no-unal --no-discordant --no-mixed --very-sensitive -I 30 -X 1000”. Alignments mapping to the mitochondrial genome were excluded from downstream analyses. PCR duplicates were identified and removed with Picard (version 2.25.7, https://broadinstitute.github.io/picard/). The BEDTools toolkit (version 2.31.1)^46^ was applied to compute reads-per-million (RPM)-normalized bedGraph coverage files, which were then converted to bigWig format using the UCSC bedGraphToBigWig tool (version 2.10)^47^. Integrative Genomics Viewer (IGV) (version 2.8.10)^48^ was used to visualize ATAC-seq coverage tracks. Peak calling for each sample was carried out with MACS2 (version 2.2.9.1)^49^, with parameters “--keep-dup all --nomodel --shift -100 --extsize 200 -q 0.01 -g mm”. Peaks overlapping genomic blacklist regions were then removed with BEDTools.

#### ATAC-seq differential peak analysis

To identify peaks that were gained or lost during the CIM induction process between 0 h and 36 h cells, differential peak analysis was performed using the following procedure as described previously^50^. Briefly, peaks called by MACS2 from both timepoints were merged using the merge subcommand from BEDTools and extended in both directions by 250 bp. Read counts for these merged regions were then quantified using the multicov subcommand. The raw counts were normalized to counts per million (CPM), and differential peaks were identified using the “exactTest” method from the edgeR package.

#### Average profile plots for ATAC-seq samples

Average profile plots (**Fig. 1e, f**) were generated using the computeMatrix and plotProfile tools from deepTools suite^51^.

#### Motif enrichment analysis

The top 50% ATAC-seq peaks (ranked by signal) from CIM-treated cells at 0 h, 12 h, 24 h and 36 h were extracted and extended ±200 bp around their summits. All peaks located within ±500 bp of TSS or overlapping CpG islands were filtered out using BEDTools. SP2 binding motif enrichment within these open chromatin regions (**Fig. 3c**) was assessed using findMotifsGenome.pl from the HOMER suite (version 5.1)^52^.

### ChIP-seq data analysis

Quality check of sequence libraries was performed by FastQC (version 0.11.8, https://www.bioinformatics.babraham.ac.uk/projects/fastqc/) with default parameters. Raw fastq reads were trimmed by trim_galore to remove any adapter sequences. The trimmed reads were aligned to the mouse reference genome mm10 by bowtie2 with options: --no-unal --no-discordant --no-mixed --very-sensitive --score-min L,0,-0.4 -X 1000. Duplicates were removed by MarkDuplicates from Picard tools. Peaks were called using MACS2 with the parameters “--keep-dup all --nomodel --shift -100 --extsize 200 -q 0.01 -g mm”. All peaks that matched the ENCODE blacklist regions of the genome were filtered using BEDTools. Peak annotation was performed using the R package ChIPseeker (version 1.26.2)^53^.

For visualization of ChIP-seq as tracks, bamtobed and genomecov from BEDTools toolkit were used to create bedGraph files consisting of RPM values. The bedGraph files were converted to bigWig files for fast query retrievals using UCSC bedGraphToBigWig. ChIP-seq coverage tracks were visualized using IGV (version 2.8.10)^48^. The BEDTools intersectBed tool was used to determine the proportion of marker genes whose promoter regions overlap SP2 peaks. The computeMatrix and plotHeatmap tools from the deepTools suite were used to generate enrichment heatmaps^51^.

## Acknowledgements

We thank Yangming Wang for providing the MERVL-reporter mES cell line. This work was financially supported by the National Key R&D Program of China (2024YFA1802300), The National Natural Science Foundation of China (32225012, 82470704), China Postdoctoral Science Foundation (2026M793072), Science and Technology Planning Project of Guangdong Province, China (2023TQ07A630, 2023B1212060050, 2023B1212120009), and Health@InnoHK Program launched by Innovation Technology Commission of the Hong Kong SAR, P. R. China.

## Author Contribution

S.C. and P.Z. initiated the project. J.C., S.C., S.H., P.Z., and H.W. designed the study. H.W. and P.Z. performed the main experiments. H.W. and S.H. interpreted the data. S.H. performed the bioinformatics analyses. H.L., H.C., S.X., E.G., B.C. and X.Q. assisted with parts of the experiments. S.H. and H.W. wrote the manuscript, and J.C., S.C. and D.P. helped to improve it. J.C., S.C., and H.W. supervised this project.

## Data availability

The data supporting the findings of this study are available from the corresponding author upon reasonable request. All the next-generation-sequencing data generated in the current study can be obtained at Genome Sequence Archive (GSA) under accession number: CRA033012.

## Supplementary information

Supplementary Table S1. Lists of primers used in this study. Supplementary Table S2. Summary of public datasets used in this study.

## Supplementary Figures

**Supplementary Fig. S1.**
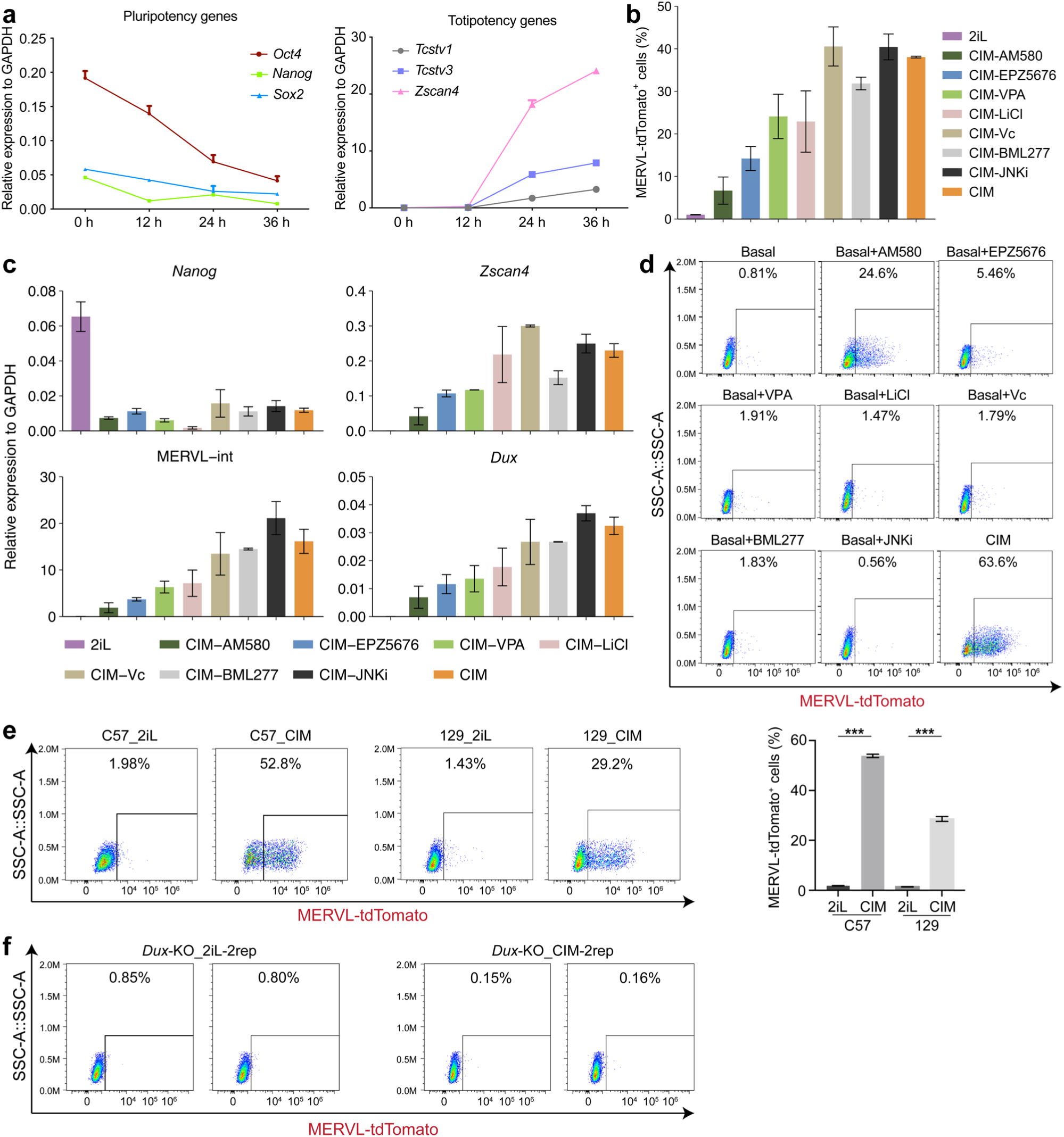
characterization of chemical induction medium, related to Fig. 1. **a**, Expression changes of pluripotency genes (*Oct4*, *Nanog*, and *Sox2*) and totipotency genes (*Tcstv1*, *Tcstv3*, and *Zscan4*) during 2C-like induction at 0 h, 12 h, 24 h and 36 h. Data are mean ± s.d., *n* = 2 independent experiments. **b**, Percentage of MERVL-tdTomato^+^ cells following culture in 2iL medium, CIM, or CIM lacking the indicated individual components. Data are mean ± s.d., *n* = 2 independent experiments. **c**, Bar plots showing the expression of *Nanog*, *Zscan4*, MERVL-int, and *Dux* in mESCs cultured in 2iL medium, CIM, or CIM lacking the indicated individual components. Data are mean ± s.d., *n* = 2 independent experiments. **d**, FACS analysis showing the percentage of MERVL-tdTomato^+^ cells in basal medium with or without the indicated compounds, or in CIM. **e**, Left panel: FACS analysis showing the percentage of MERVL-tdTomato^+^ cells tested on C57 and 129 mESCs with CIM. Right panel: Bar plot showing the quantification of the corresponding FACS analysis. Shown are mean ± s.d., *n* = 3 independent experiments. \*\*\**P* < 0.001; two-tailed unpaired Student’s *t*-test. **f**, FACS analysis showing the percentage of MERVL-tdTomato^+^ cells in *Dux*-knockout (KO) mESCs cultured in 2iL medium or CIM. Two independent experiments are shown.

**Supplementary Fig. S2.**
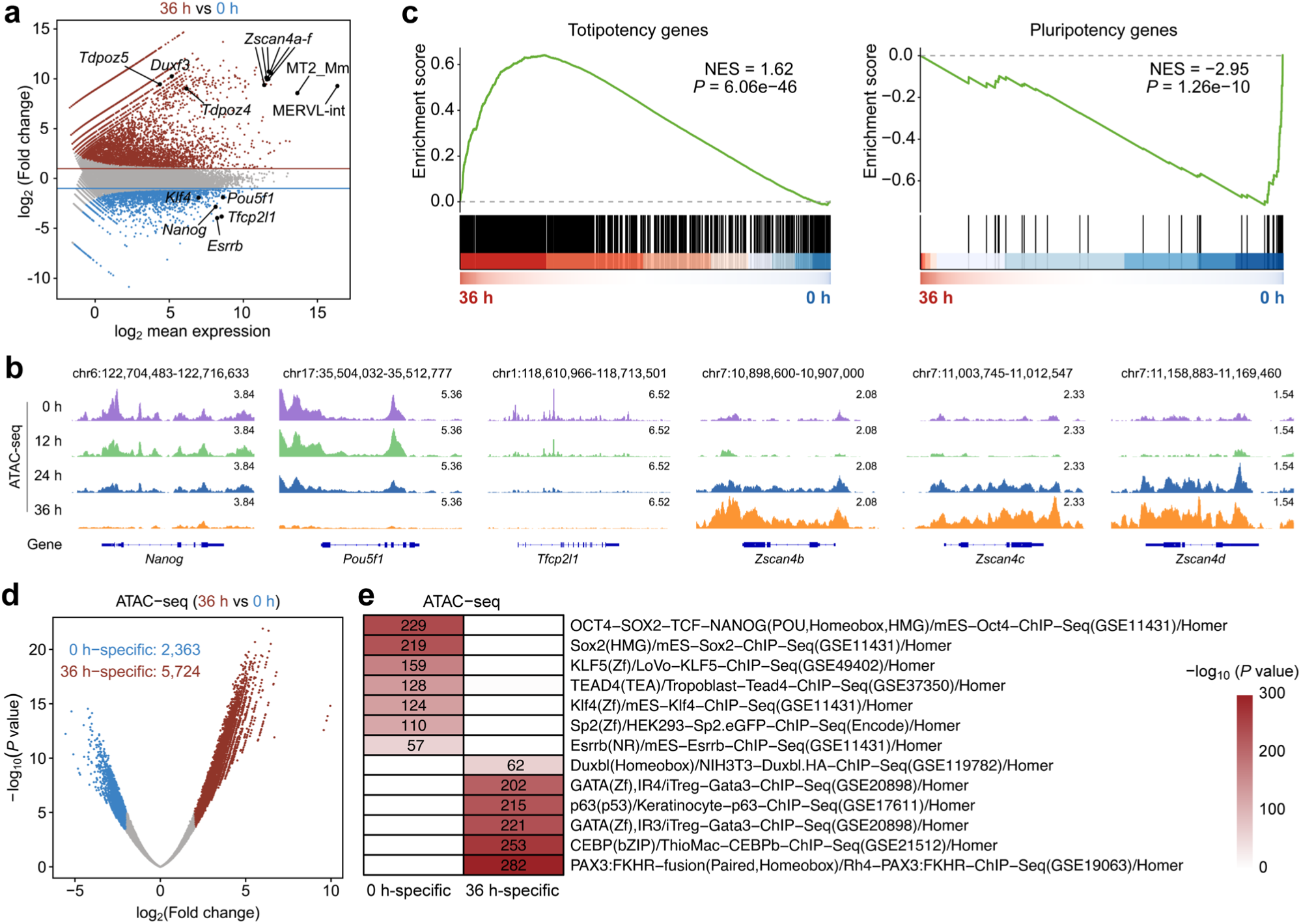
The transcriptomic and epigenomic features of CIM-induced cells are similar to 2-cell blastomeres, related to Fig. 1. **a**, MA plot showing differentially expressed genes and TE subfamilies between 36 h and 0 h during chemical reprogramming. Red points: upregulated (log_2_FC ≥ 1, *P* < 0.01); Blue points: downregulated (log_2_FC ≤ -1, *P* < 0.01). **b**, Integrative Genomics Viewer (IGV) snapshots displaying ATAC-seq signals around representative totipotency and pluripotency genes. **c**, Gene set enrichment analysis (GSEA) of totipotency genes (left) and pluripotency genes (right) in mESCs treated with CIM for 36 h or 0 h. **d**, Volcano plot showing the ATAC-seq differential peaks between 36 h- and 0 h-chemical induction. Blue, gray, and red indicate lost (0 h-specific; log_2_FC ≤ -2, *P* < 0.01), stable, and gained (36 h-specific; log_2_FC ≥ 2, *P* < 0.01) sites, respectively. **e**, Heatmap of different transcription factors’ motif analysis in the 0 h-specific and 36 h-specific peak clusters identified in (**d**).

**Supplementary Fig. S3.**
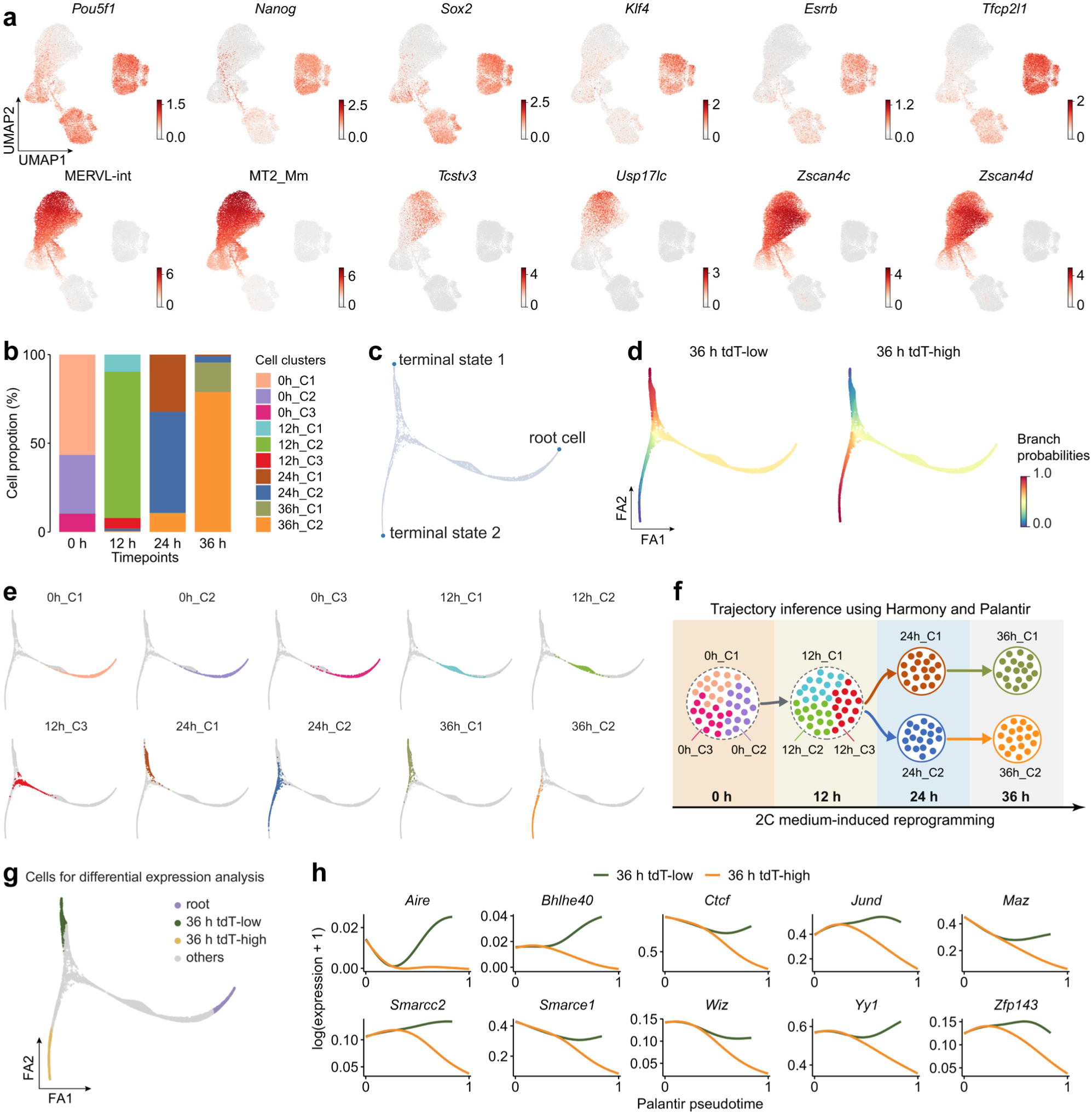
Characterization of branched trajectory in chemical reprogramming using scRNA-seq, related to Fig. 2. **a**, UMAP plot showing the expression of the indicated genes or TEs during chemical reprogramming. Top, pluripotency markers; Bottom, totipotency markers. **b**, Stacked bar plot showing the proportions of the ten cell clusters at different timepoints during chemical reprogramming. **c**, The root cell and terminal states selected for pseudotime inference. **d**, ForceAtlas2 layout showing branch probabilities for the 36 h tdT-low and 36 h tdT-high branches. **e**, Trajectory graph stratified by ten cell clusters, with each panel depicting the trajectory of a single cell cluster, extracted from the middle panel of Fig. 2d. **f**, Palantir-based schematic of cluster transitions. Colored dots represent cell clusters, and directed arrows indicate inferred cell fate transitions. **g**, ForceAtlas2 layout of the reprogramming trajectories, highlighting three cell groups: root cells (500 cells with the lowest pseudotime), and terminal populations from the 36 h tdT-low and 36 h tdT-high branches (500 cells each with the largest pseudotime). Cells are colored by group identity. **h**, Gene expression trends along Palantir pseudotime for chromatin architectural proteins and other chromatin regulators.

**Supplementary Fig. S4.**
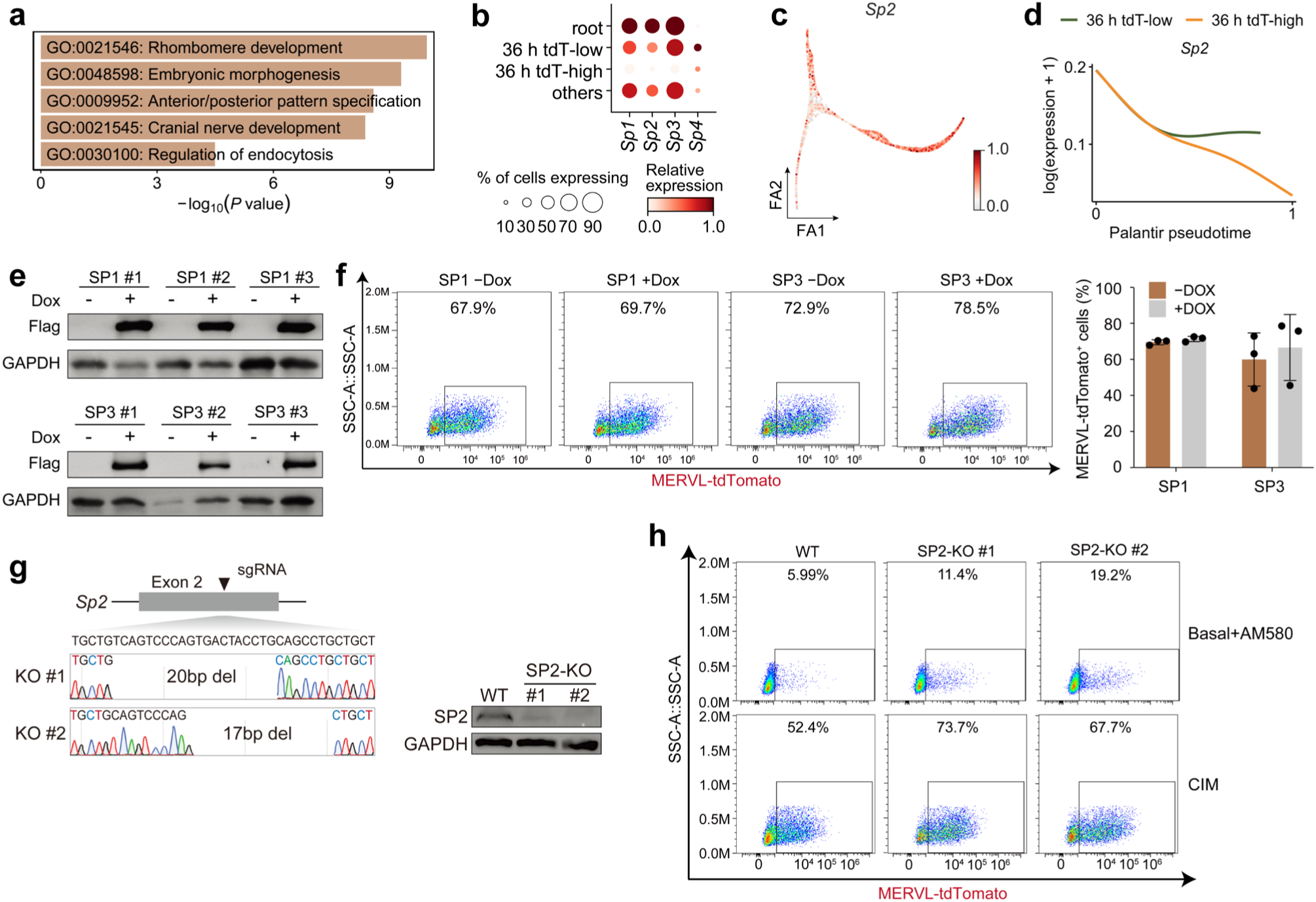
Expression and functional analyses of SP family members during 2C-like reprogramming, related to Fig. 3. **a**, Metascape bar plot showing enriched pathways among 36h_C1 marker genes predicted by RcisTarget to be targets of SP family transcription factors. **b**, Dot plot showing the expression pattern of *Sp* family genes in the indicated cell groups. The dot size represents the percentage of cells in each cluster expressing the corresponding genes, and the color intensity reflects the scaled expression values. **c**, ForceAtlas2 layout showing the expression levels of *Sp2* in different branches. **d**, Comparison of gene expression trends along Palantir pseudotime for *Sp2*. **e**, Western blot showing SP1 and SP3 protein levels in 2iL medium with or without Dox. **f**, Left panel: FACS analysis showing the percentage of MERVL-tdTomato^+^ cells of SP1 and SP3 over-expressing mESC lines in CIM with or without Dox (*n* = 1). Right panel: Bar plot showing the quantification of the corresponding FACS analysis. Shown are mean ± s.d., *n* = 3 independent experiments. **g**, Sanger sequencing analysis showing the missing nucleotides induced by CRISPR-Cas9 at exon 2 of *Sp2* gene on the left, and the western blot result showing the deletion of SP2 on the protein level on the right. **h**, Representative FACS plots showing the percentage of MERVL-tdTomato^+^ cells in WT and two SP2-KO mESC clones cultured in basal medium supplemented with AM580 or in CIM.

**Supplementary Fig. S5.**
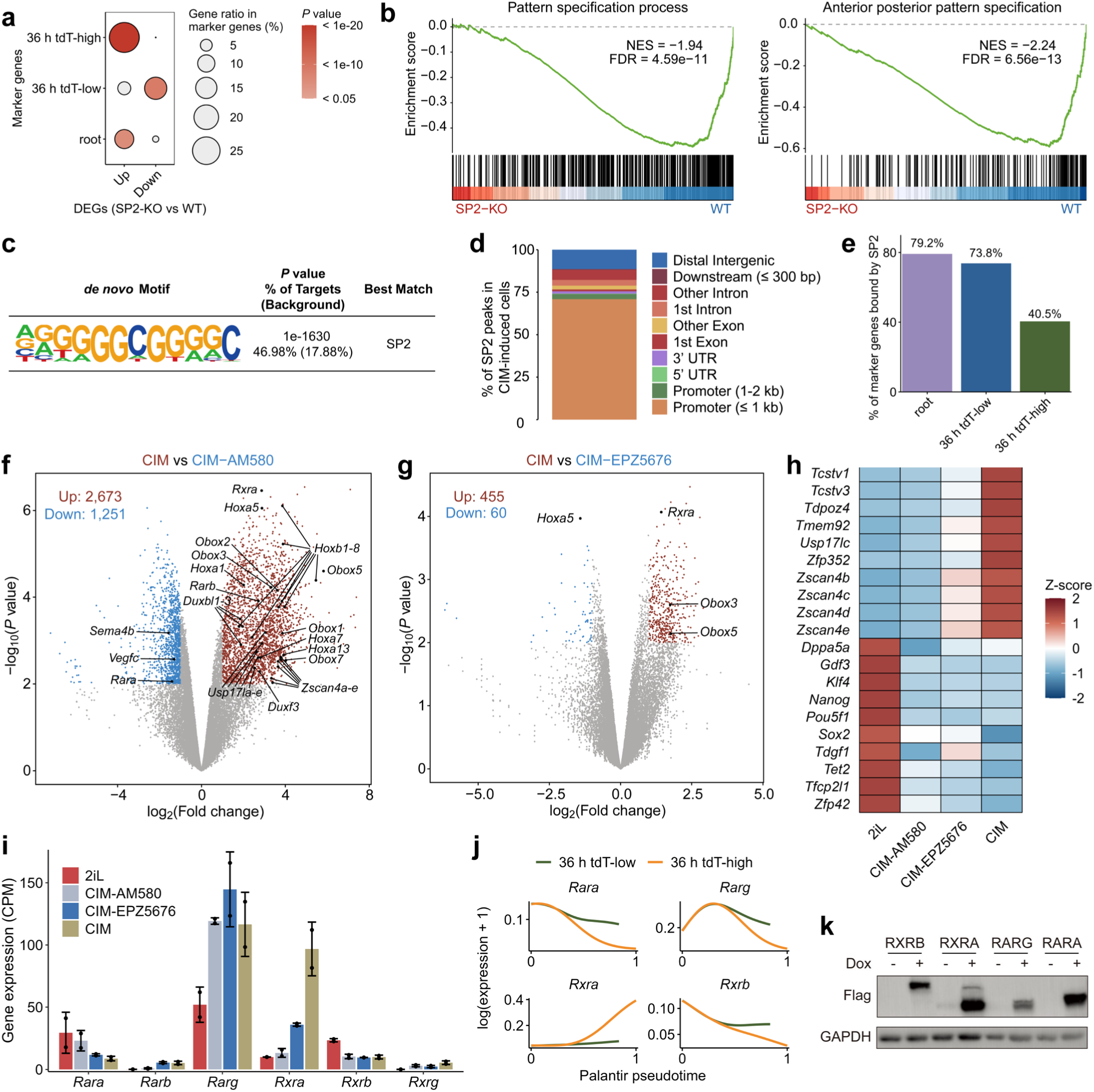
SP2 regulates *Hox* gene expression in an RA-dependent manner, related to Fig. 4. **a**, Comparison of the enrichment of genes up- or downregulated after SP2 deletion among marker genes of the root, 36 h tdT-low and 36 h tdT-high cell groups. Point size represents the gene ratio of marker genes, and the red gradient represents the *P* values. *P* values were calculated by a one-tailed hypergeometric test. **b**, GSEA plots of the Pattern specification process (left) and Anterior/posterior pattern specification (right) gene sets comparing SP2-KO and WT mESCs cultured in CIM for 36 h. **c**, HOMER *de novo* motif search for the SP2 ChIP-seq peaks identified the SP2 motif as the top hit. **d**, Bar plot showing the genomic distribution of SP2 peaks. **e**, Bar plots showing the percentage of marker genes from each cell group that are bound by SP2. **f and g**, Volcano plots showing differentially expressed genes between CIM and CIM lacking AM580 (**f**) or EPZ5676 (**g**). Red points: upregulated (log_2_FC ≥ 1, *P* < 0.01); Blue points: downregulated (log_2_FC ≤ -1, *P* < 0.01). **h**, Heatmap showing the relative expression of totipotency and pluripotency genes in mESCs cultured in 2iL medium, CIM without AM580, CIM without EPZ5676, or CIM. **i**, Bar plots showing the expression levels of RA receptors in mESCs cultured in 2iL medium, CIM without AM580, CIM without EPZ5676, or CIM. Shown are mean ± s.d., *n* = 2 independent experiments. **j**, Gene expression trends of RA receptors, including *Rara*, *Rarg*, *Rxra*, and *Rxrb*, along Palantir pseudotime in the 36 h tdT-low and 36 h tdT-high branches. **k**, Western blot analysis confirming Dox-induced overexpression of RA receptors.

## Reference

1. Tarkowski, A.K. Experiments on the development of isolated blastomers of mouse eggs. Nature 184, 1286–1287 (1959).

2. Posfai, E. et al. Evaluating totipotency using criteria of increasing stringency. Nat Cell Biol 23, 49–60 (2021).

3. Cockburn, K. & Rossant, J. Making the blastocyst: lessons from the mouse. J. Clin. Invest. 120, 995–1003 (2010).

4. Lu, F. & Zhang, Y. Cell totipotency: molecular features, induction, and maintenance. Natl Sci Rev 2, 217–225 (2015).

5. Macfarlan, T.S. et al. Embryonic stem cell potency fluctuates with endogenous retrovirus activity. Nature 487, 57–63 (2012).

6. Rodriguez-Terrones, D. et al. A molecular roadmap for the emergence of early-embryonic-like cells in culture. Nat. Genet. 50, 106–119 (2018).

7. Hendrickson, P.G. et al. Conserved roles of mouse DUX and human DUX4 in activating cleavage-stage genes and MERVL/HERVL retrotransposons. Nat. Genet. 49, 925–934 (2017).

8. Wen, X., Lin, Z., Wu, H., Cao, L. & Fu, X. Zfp281 Inhibits the Pluripotent-to-Totipotent State Transition in Mouse Embryonic Stem Cells. Front Cell Dev Biol 10, 879428 (2022).

9. Fu, X., Wu, X., Djekidel, M.N. & Zhang, Y. Myc and Dnmt1 impede the pluripotent to totipotent state transition in embryonic stem cells. Nat Cell Biol 21, 835–844 (2019).

10. Wu, K. et al. SETDB1-Mediated Cell Fate Transition between 2C-Like and Pluripotent States. Cell Rep. 30, 25–36.e26 (2020).

11. Liu, J. et al. The RNA m(6)A reader YTHDC1 silences retrotransposons and guards ES cell identity. Nature 591, 322–326 (2021).

12. Chen, C. et al. Nuclear m(6)A reader YTHDC1 regulates the scaffold function of LINE1 RNA in mouse ESCs and early embryos. Protein Cell 12, 455–474 (2021).

13. Olbrich, T. et al. CTCF is a barrier for 2C-like reprogramming. Nat. Commun. 12, 4856 (2021).

14. Li, P. et al. Rif1 promotes a repressive chromatin state to safeguard against endogenous retrovirus activation. Nucleic Acids Res 45, 12723–12738 (2017).

15. Tagliaferri, D. et al. Retinoic Acid Induces Embryonic Stem Cells (ESCs) Transition to 2 Cell-Like State Through a Coordinated Expression of Dux and Duxbl1. Front Cell Dev Biol 7, 385 (2019).

16. Hu, Y. et al. Induction of mouse totipotent stem cells by a defined chemical cocktail. Nature 617, 792–797 (2023).

17. Xu, Y. et al. Derivation of totipotent-like stem cells with blastocyst-like structure forming potential. Cell Res. 32, 513–529 (2022).

18. Li, H. et al. A complete model of mouse embryogenesis through organogenesis enabled by chemically induced embryo founder cells. Cell 188, 5912–5930.e5920 (2025).

19. Chen, J. et al. Rational optimization of reprogramming culture conditions for the generation of induced pluripotent stem cells with ultra-high efficiency and fast kinetics. Cell Res. 21, 884–894 (2011).

20. Cao, S. et al. Chromatin Accessibility Dynamics during Chemical Induction of Pluripotency. Cell Stem Cell 22, 529–542.e525 (2018).

21. Yang, J. et al. Establishment of mouse expanded potential stem cells. Nature 550, 393–397 (2017).

22. Chen, X. et al. A fast chemical reprogramming system promotes cell identity transition through a diapause-like state. Nat Cell Biol 25, 1146–1156 (2023).

23. Shen, H. et al. Mouse totipotent stem cells captured and maintained through spliceosomal repression. Cell 184, 2843–2859.e2820 (2021).

24. Yang, M. et al. Chemical-induced chromatin remodeling reprograms mouse ESCs to totipotent-like stem cells. Cell Stem Cell 29, 400–418.e413 (2022).

25. Deng, Q., Ramsköld, D., Reinius, B. & Sandberg, R. Single-cell RNA-seq reveals dynamic, random monoallelic gene expression in mammalian cells. Science 343, 193–196 (2014).

26. Shi, P. et al. USP17L promotes the 2-cell-like program through deubiquitination of H2AK119ub1 and ZSCAN4. Nat. Commun. 16, 7071 (2025).

27. Zhu, Y. et al. Relaxed 3D genome conformation facilitates the pluripotent to totipotent-like state transition in embryonic stem cells. Nucleic Acids Res 49, 12167–12177 (2021).

28. Iturbide, A. et al. Retinoic acid signaling is critical during the totipotency window in early mammalian development. Nat. Struct. Mol. Biol. 28, 521–532 (2021).

29. Da, C.M. et al. Transcription Factor SP2 Regulates Ski-mediated Astrocyte Proliferation In Vitro. Neuroscience 479, 22–34 (2021).

30. Zhu, Y. et al. Sp2 promotes invasion and metastasis of hepatocellular carcinoma by targeting TRIB3 protein. Cancer Med 9, 3592–3603 (2020).

31. Phan, D. et al. Identification of Sp2 as a transcriptional repressor of carcinoembryonic antigen-related cell adhesion molecule 1 in tumorigenesis. Cancer Res. 64, 3072–3078 (2004).

32. Johnson, C.A. & Ghashghaei, H.T. Sp2 regulates late neurogenic but not early expansive divisions of neural stem cells underlying population growth in the mouse cortex. Development 147 (2020).

33. Dobin, A. et al. STAR: ultrafast universal RNA-seq aligner. Bioinformatics 29, 15–21 (2013).

34. Jin, Y., Tam, O.H., Paniagua, E. & Hammell, M. TEtranscripts: a package for including transposable elements in differential expression analysis of RNA-seq datasets. Bioinformatics 31, 3593–3599 (2015).

35. Robinson, M.D. & Oshlack, A. A scaling normalization method for differential expression analysis of RNA-seq data. Genome Biol 11, R25 (2010).

36. Robinson, M.D., McCarthy, D.J. & Smyth, G.K. edgeR: a Bioconductor package for differential expression analysis of digital gene expression data. Bioinformatics 26, 139–140 (2010).

37. Korotkevich, G., Sukhov, V. & Sergushichev, A. Fast gene set enrichment analysis. bioRxiv, 060012 (2019).

38. He, J. et al. Identifying transposable element expression dynamics and heterogeneity during development at the single-cell level with a processing pipeline scTE. Nat. Commun. 12, 1456 (2021).

39. Wolock, S.L., Lopez, R. & Klein, A.M. Scrublet: Computational Identification of Cell Doublets in Single-Cell Transcriptomic Data. Cell Syst 8, 281–291.e289 (2019).

40. Aibar, S. et al. SCENIC: single-cell regulatory network inference and clustering. Nat. Methods 14, 1083–1086 (2017).

41. Nowotschin, S. et al. The emergent landscape of the mouse gut endoderm at single-cell resolution. Nature 569, 361–367 (2019).

42. Setty, M. et al. Characterization of cell fate probabilities in single-cell data with Palantir. Nat. Biotechnol. 37, 451–460 (2019).

43. Petitpré, C. et al. Single-cell RNA-sequencing analysis of the developing mouse inner ear identifies molecular logic of auditory neuron diversification. Nat. Commun. 13, 3878 (2022).

44. Zhou, Y. et al. Metascape provides a biologist-oriented resource for the analysis of systems-level datasets. Nat. Commun. 10, 1523 (2019).

45. Langmead, B. & Salzberg, S.L. Fast gapped-read alignment with Bowtie 2. Nat. Methods 9, 357–359 (2012).

46. Quinlan, A.R. & Hall, I.M. BEDTools: a flexible suite of utilities for comparing genomic features. Bioinformatics 26, 841–842 (2010).

47. Kuhn, R.M., Haussler, D. & Kent, W.J. The UCSC genome browser and associated tools. Brief. Bioinform. 14, 144–161 (2013).

48. Robinson, J.T. et al. Integrative genomics viewer. Nat. Biotechnol. 29, 24–26 (2011).

49. Zhang, Y. et al. Model-based analysis of ChIP-Seq (MACS). Genome Biol 9, R137 (2008).

50. Peng, B., Wang, Q., Zhang, F., Shen, H. & Du, P. Mouse totipotent blastomere-like cells model embryogenesis from zygotic genome activation to post implantation. Cell Stem Cell 32, 391–408.e311 (2025).

51. Ramírez, F., Dündar, F., Diehl, S., Grüning, B.A. & Manke, T. deepTools: a flexible platform for exploring deep-sequencing data. Nucleic Acids Res 42, W187–191 (2014).

52. Heinz, S. et al. Simple combinations of lineage-determining transcription factors prime cis-regulatory elements required for macrophage and B cell identities. Mol Cell 38, 576–589 (2010).

53. Yu, G., Wang, L.G. & He, Q.Y. ChIPseeker: an R/Bioconductor package for ChIP peak annotation, comparison and visualization. Bioinformatics 31, 2382–2383 (2015).

