## Supplementary Tables for "SP2 impedes the pluripotent-to-totipotent state transition in an RA-dependent manner"

**Supplementary Table 1. Lists of primers used in this study.**

| Primers | Sequences |
|---|---|
| Gapdh-F | ACCTGCCAAGTATGATGAC |
| Gapdh-R | GGGAGTTGCTGTTGAAGT |
| Oct4-F | CATTGAGAACCGTGTGAG |
| Oct4-R | TGAGTGATCTGCTGTAGG |
| Nanog-F | CTCAAGTCCTGAGGCTGACA |
| Nanog-R | TGAAACCTGTCCTTGAGTGC |
| Zscan4-F | GTCCTGACAGAGGCCTGCC |
| Zscan4-R | GAGATGTCTGAAGAGGCAAT |
| Tcstv1-F | GGCACAGAGGTCTTCCAGAG |
| Tcstv1-R | CAGACCTTTCTCCCTGAGCA |
| Tcstv3-F | AGAAAGGGCTGGAACTTGTGACCT |
| Tcstv3-R | AAAGCTCTTTGAAGCCATGCCCAG |
| Mervl-LTR-F | CTTCCATTCACAGCTGCGACTG |
| Mervl-LTR-R | CTAGAACCACTCCTGGTACCAAC |

**Supplementary Table 2. Summary of public datasets used in this study.**

| Data type | Sample | NCBI GEO / EMBL-EBI ArrayExpress accession |
|---|---|---|
| ATAC-seq | 2-cell rep1 | GSM1933924 |
| | 2-cell rep2 | GSM1933925 |
| | 4-cell rep1 | GSM1625847 |
| | 4-cell rep2 | GSM1933927 |
| | 8-cell rep1 | GSM1933928 |
| | 8-cell rep2 | GSM1933929 |
| | ICM rep1 | GSM1933930 |
| | ICM rep2 | GSM2156963 |
| RNA-seq | TBLC-P6 | GSM5160100 |
| | TLSC-C57-P12 | GSM5603639 |
| | TLSC-2C-P8 | GSM5603636 |
| | TLSC-2C-P14 | GSM5603637 |
| | mESC-C57 | GSM5603641 |
| | 2CLC_rep1 | GSM5065808 |
| | 2CLC_rep2 | GSM5065809 |
| | TPSC#1_rep1 | GSM5558728 |
| | TPSC#1_rep2 | GSM5558729 |
| | TPSC#2_rep1 | GSM5558730 |
| | TPSC#2_rep2 | GSM5558731 |
| | TPSC#3_rep1 | GSM5558732 |
| | TPSC#3_rep2 | GSM5558733 |
| | D-EPSC#1_rep1 | GSM5558734 |
| | D-EPSC#1_rep2 | GSM5558735 |
| | D-EPSC#2_rep1 | GSM5558736 |
| | D-EPSC#2_rep2 | GSM5558737 |
| | D-EPSC#3_rep1 | GSM5558738 |
| | D-EPSC#3_rep2 | GSM5558739 |
| | ciTotiSC-P1_rep1 | GSM5603033 |
| | ciTotiSC-P1_rep2 | GSM5603034 |
| | ciTotiSC-P2_rep1 | GSM5603035 |
| | ciTotiSC-P2_rep2 | GSM5603036 |
| | ciTotiSC-P4_rep1 | GSM5603037 |
| | ciTotiSC-P4_rep2 | GSM5603038 |
| | ciTotiSC-P8_rep1 | GSM5603039 |
| | ciTotiSC-P8_rep2 | GSM5603040 |
| | OG2_mESC_rep1 | GSM5603041 |
| | OG2_mESC_rep2 | GSM5603042 |
| | mESC rep1 | GSM1625873 |
| | mESC rep2 | GSM1625874 |
| | D0_2C-_rep1 | GSM3436706 |
| | D0_2C-_rep2 | GSM3436707 |

| | | |
|---|---|---|
| | D1_2C-_rep1 | GSM3436708 |
| | D1_2C-_rep2 | GSM3436709 |
| | D1_2C+_rep1 | GSM3436710 |
| | D1_2C+_rep2 | GSM3436711 |
| | mESC_rep1 | GSM4835983 |
| | mESC_rep2 | GSM4835984 |
| | mESC_rep3 | GSM4835985 |
| | mESC_rep4 | GSM4835986 |
| | 2CLC_rep1 | GSM4835987 |
| | 2CLC_rep2 | GSM4835988 |
| | 2CLC_rep3 | GSM4835989 |
| | 2CLC_rep4 | GSM4835990 |
| single-cell RNA-seq | SC-TBLC | GSM5195025 |
| | ciTotiSC | GSM5603043 |
| | zygote | GSE45719 |
| | early2cell | |
| | mid2cell | |
| | late2cell | |
| | C57twocell | |
| | 4cell | |
| | 8cell | |
| | 16cell | |
| | earlyblast | |
| | midblast | |
| | lateblast | |
| | mESC RA 0 h | E-MTAB-8869 |
| | mESC RA 2 h | |
| | mESC RA 12 h | |
| | mESC RA 48 h | |
